# Synthetic niches enable co-culture bioprocessing but are prone to mutational escape

**DOI:** 10.1101/2025.11.24.690348

**Authors:** Vincent Vandenbroucke, Juan Andrés Martínez, Lucas Henrion, Andrew Zicler, Samuel Telek, Laurie Josselin, Frank Delvigne

## Abstract

Stabilizing microbial co-cultures is a central challenge for bioproduction. While division of labor between strains can enhance efficiency, it often results in population instability over time. Classical strategies, including cross-feeding, quorum sensing, and toxin-antitoxin modules, often rely on complex ecological interactions that are difficult to predict or maintain under bioprocess conditions. Here, we introduce synthetic niches as an alternative framework, using genetic toggle switches that couple growth to defined phenotypic states. We engineered two auxotrophic strains, TOGGLE_green and TOGGLE_yellow, in which growth is linked to either GFP- or YFP-expressing states and assessed their behavior in continuous bioreactor cultures using automated flow cytometry. Unexpectedly, the introduction of auxotrophic pressure reshaped circuit function i.e., instead of maintaining bistability, toggle strains behaved as unidirectional inducible systems that reverted upon inducer withdrawal. This feature enabled simplified control with a single input but also revealed a vulnerability to mutational escape under intensified cultivation. A simple repression-based ODE model recapitulated the reversible dynamics, but deviations under prolonged operation highlighted the rapid evolutionary erosion of control. Our findings demonstrate both the potential and the limitations of synthetic niches for co-culture engineering and emphasize the need to integrate evolutionary robustness into the design of next-generation bioprocess control strategies.

## Introduction

Microbial cultures provide a versatile platform for biotechnology, supporting the production of fuels, chemicals, and therapeutics. However, monocultures are often limited by metabolic burden, competing intracellular reactions, and the complexity of balancing multiple enzymatic steps within a single host [1–6]. To address these constraints, microbial co-cultures have emerged as a promising alternative. By dividing labor between strains, co-cultures reduce burden, expand the accessible metabolic space, and allow for modular pathway design [7–9]. Yet, they also introduce new challenges: co-cultures are inherently unstable, as strain ratios drift over time, frequently leading to the dominance of one of the strain and the loss of production [2,4,6]. Concepts from microbial ecology have also been applied to co-culture control, including metabolic niche theory and competitive exclusion [10–12]. These approaches are effective only when temporal dynamics are satisfied [8,13], as niche expansion or contraction often compromises long-term stability [10,14].

When natural niches and interactions are not possible, synthetic biology has been used to create them. These synthetic interactions include cross-feeding [13,15,16], quorum sensing [17], toxin-antitoxin systems and regulated killing modules [18–20]. While effective in certain contexts, these approaches often require tailoring interactions to specific pathways or strains, reducing modularity and making generalizable design difficult [16,17,20–22]. Alternatively, synthetic niches create metabolic spaces where microbes do not compete, usually via separation of carbon sources, and often require extensive genetic engineering that may adversely affect production pathways. As a result, focus has turned to creating synthetic niches that interfere minimally with normal cell metabolism and potential production pathways [18]. In this context, toggle switch circuits are particularly attractive: by coupling strain fitness to a bistable regulatory state, they enable orthogonal control of strain ratios while leaving metabolic pathways largely unaffected. Unlike cross-feeding or quorum sensing, this strategy minimizes energetic demand by relying on non-metabolizable inducers rather than costly metabolic exchanges. In this way, synthetic niches provide modularity and selectivity without constraining the underlying production pathway [1,18,21,23].

Despite their promise, most studies of synthetic consortia have been conducted under simplified cultivation conditions, such as batch or serial transfer cultures lasting fewer than 20 generations [3–6,21]. Even continuous cultures often remain below 40 generations, and typically in small-volume or microfluidic devices [18,22,24]. For instance, Sommer and colleagues observed mutations after ∼40 generations in 200 mL batch cultures [25], while Khammash and colleagues reported genetic erosion after ∼80 generations in very small-volume consortia [18]. Only a few studies extend beyond 100 generations e.g., auxotrophic cross-feeding in *E. coli* [16] or phenol degradation consortia [26] and they are largely ecological rather than production-focused. By contrast, industrial bioprocesses routinely impose more than 60 generations under high-density, large-volume conditions, where the probability of mutational escape scales with total cell divisions [25].

A further limitation lies in methods of observation. Most studies rely on manual sampling and coarse readouts such as CFU counts, DNA quantification, or bulk fluorescence [21]. A few notable exceptions, such as fluorescence-based ratio tracking in microreactors [18,23], optogenetic cybergenetic control [18,27], or microchemostat platforms [22,24], achieved higher resolution but remained restricted to small cell populations or short durations. Understanding the long-term behavior of synthetic niches under industrially relevant conditions at single-cell resolution remains an open challenge. Overcoming this limitation requires continuous, single cell monitoring tools compatible with bioreactor operation. The Segregostat platform, a cell-machine interface relying on reactive flow cytometry for real-time population monitoring and control, addresses this need by providing real-time, high-resolution feedback on strain ratios during extended cultivation [13,28].

In this work, we implement synthetic niches based on toggle switch circuits under intensified bioreactor conditions. Using auxotrophic strains and reactive flow cytometry (Segregostat) [29], we explore how auxotrophic pressure reshapes toggle dynamics over prolonged cultivation (> 80 generations). We show that while synthetic niches can achieve selective and simplified control of co-culture composition, they are also prone to mutational escape, highlighting evolutionary robustness as a key requirement for next-generation co-culture design.

## Results

### 1. Synthetic niches enable the selective control of the growth rate of microbial species involved in co-cultures

The synthetic niches approach was implemented by combining genetic toggle switches and an auxotrophy to serine via the *serA* gene, which can be deleted from the genome and used for plasmid maintenance [30]. The switches direct *serA* production for the cells and provide a growing, *serA* producing, and a non-growing, *serA* starved phenotypes. In the TOGGLE_green strain, the growth-associated phenotype was linked to the GFP-expressing state, repressed by IPTG and inducible by tetracycline (Tc). Conversely, in the TOGGLE_yellow strain, the growth-associated phenotype was coupled to the YFP-expressing state, which was inducible by IPTG and repressed by Tc (**Figure 1A**). Using these genetic constructs, the relative abundance of each strain in the bioreactor could be independently controlled through the periodic addition of IPTG or Tc (**Figure 1B**). The timing of inducer addition was guided by a previously developed cell-machine interface, the Segregostat, which relies on reactive flow cytometry for real-time population monitoring and control (**Figure 1C**) [13,29].

**Figure 1.**
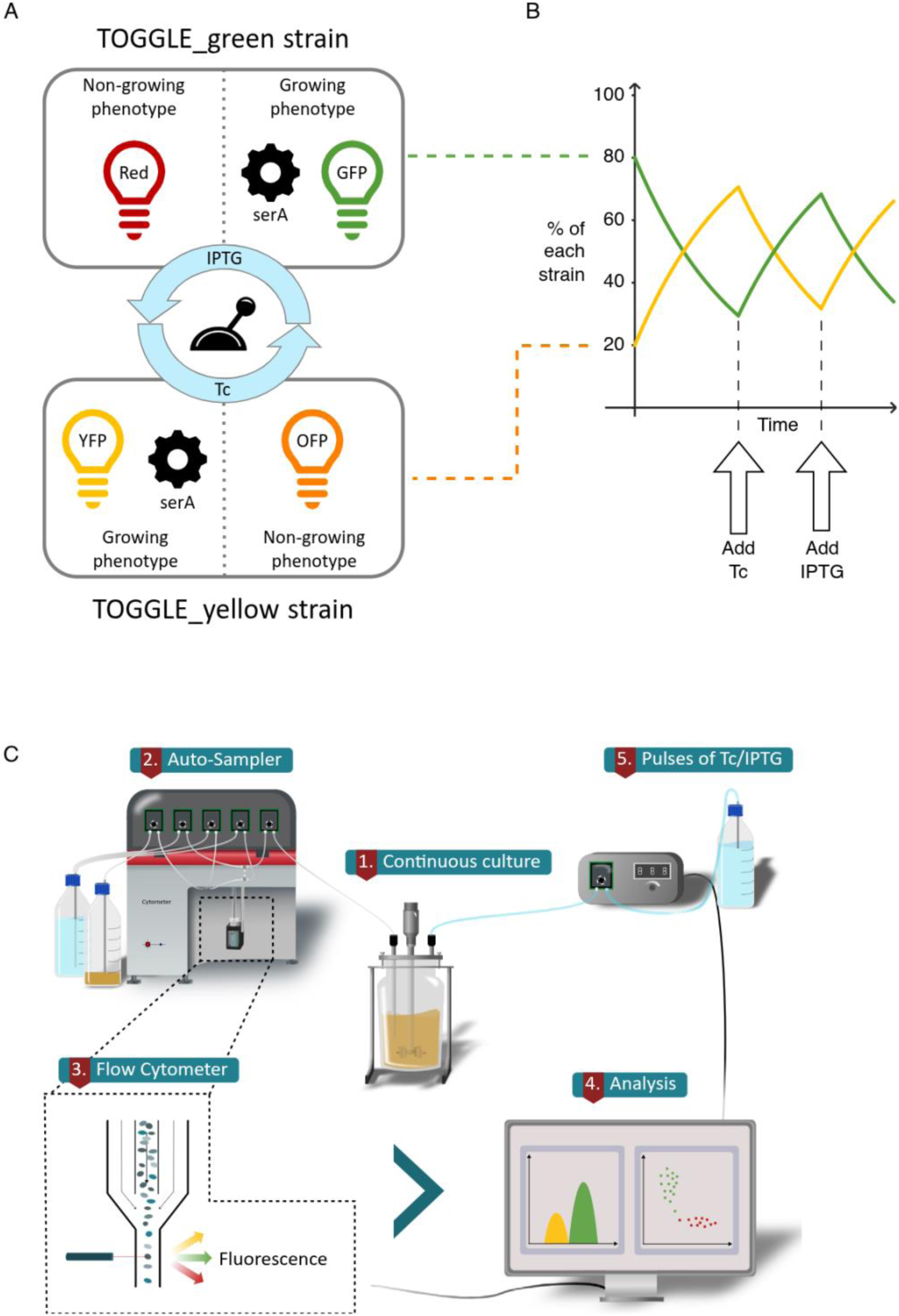
Conceptual design for implementing synthetic niches to control co-culture composition. **A** Two strains engineered with growth-dependent genetic toggle switches, where one state of the switch specifically regulates serine biosynthesis in serine auxotrophs. **B** Periodic addition of inducers enables differential control of strain growth, allowing real-time manipulation of co-culture composition. **C** A cell-machine interface (Segregostat) provides feedback-driven inducer addition for optimal co-culture regulation.

The TOGGLE_green and TOGGLE_yellow strains were constructed based on the pECJ3 plasmid, which encodes a toggle switch (**Figure 2A**) [31]. In the TOGGLE_green strain, the pECJ3 backbone was retained, and the GFP state was complemented with the *serA* gene, enabling serine biosynthesis upon induction (**Figure 2B**). Microplate cultivation revealed a marked growth difference between the GFP and mCherry states of the toggle, corresponding to a 44% decrease in growth rate (**Figure 2C**).

**Figure 2.**
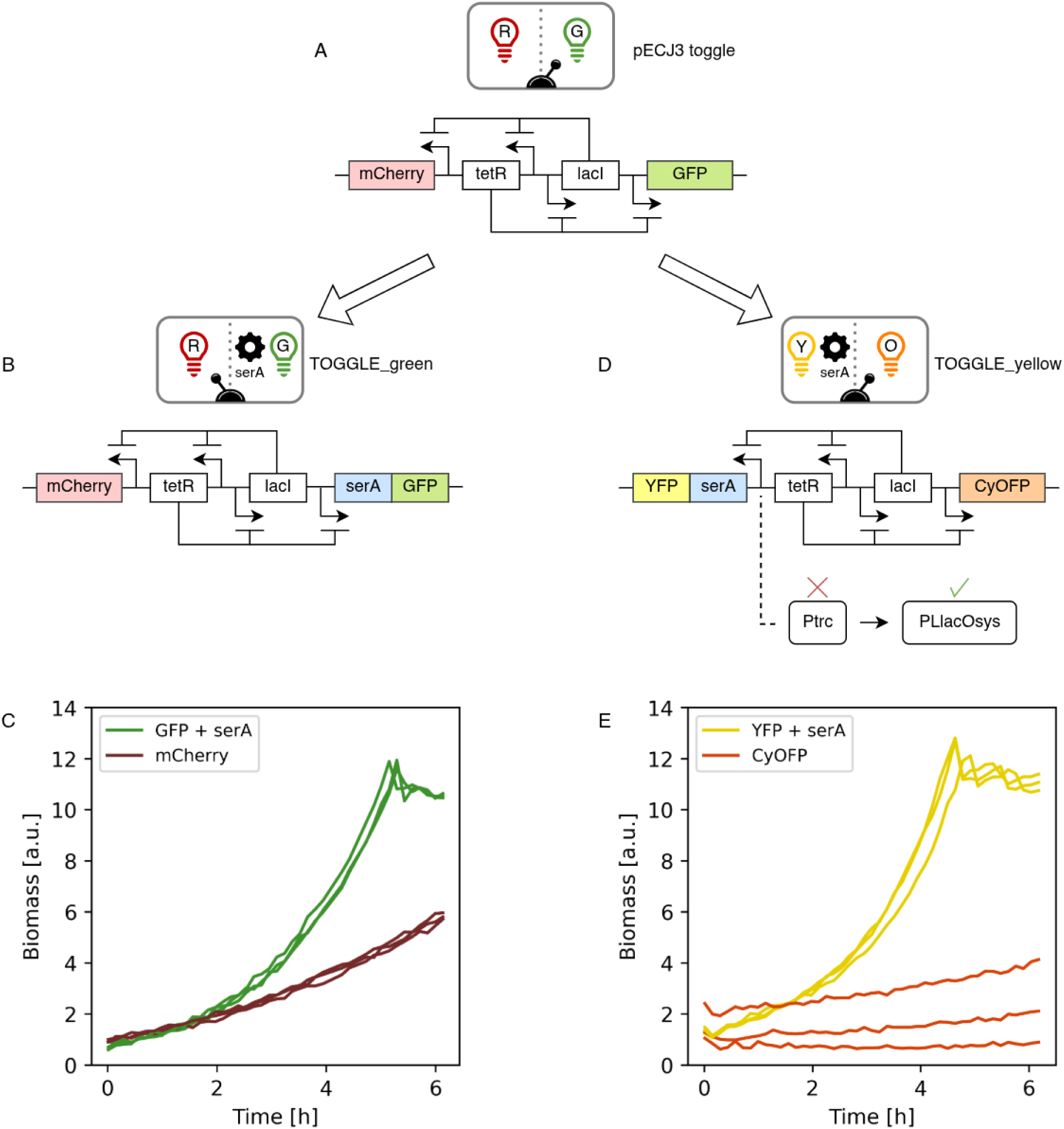
Toggle switch design for implementing synthetic niches. **A** Genetic toggle switch architecture carried on the pECJ3 backbone plasmid, used in both strains. **B** Schematic representation of the TOGGLE_green construct showing the main regulatory elements. **C** Biomass dynamics of TOGGLE_green flipped to either the GFP or mCherry state (*n* = 3; 0.53 h^-1^; σ=0.015 h^-1^ and 0.29 h^-1^; σ=0.010 h^-1^; p=3E-5). **D** Schematic representation of the TOGGLE_yellow construct showing the main regulatory elements. **E** Biomass dynamics of TOGGLE_yellow flipped to either the YFP or CyOFP state (*n* = 3; 0.44 h^-1^; σ=0.021 h^-1^ and 0.26 h^-1^; σ=0.054 h^-1^; p=8E-3).

For the TOGGLE_yellow strain, mCherry and GFP were replaced by sYFP2 (YFP) and CyOFP1 (CyOFP), respectively, with the YFP state coupled to *serA* expression and growth (**Figure 2D**). The initial construct, however, showed no significant growth difference between the two states due to leakiness of the P_trc_ promoter driving YFP and *serA*. Replacing this promoter with the tighter P_LlacOsys_ promoter restored toggle functionality, yielding a comparable 42% growth difference between the two states (**Figure 2E**). P_LlacOsys_ was derived from the P_LlacO_ promoter, which is tighter than P_trc_, and the lacOsys binding site, which enhances LacI binding (see **Materials and Methods**) [32–34]. Each TOGGLE system was engineered with two distinct fluorescent proteins to mark the alternative states. For subsequent analyses, we focused on the *serA*-associated GFP in TOGGLE_green and YFP in TOGGLE_yellow, as these provide a direct proxy for the growth status of individual cells. In contrast, CyOFP and mCherry served primarily as confirmation that the two switch states were mutually exclusive but offered little additional information.

### 2. Auxotrophic pressure induces cell escape from the toggle switch

To validate the predictive capacity of the pECJ3 toggle, the system was first evaluated in continuous culture with flow cytometry monitoring. Subsequently, the derived TOGGLE construct, built from pECJ3, was tested under similar conditions to assess whether it retained similar dynamics. However, unexpected population dynamics emerged upon introducing auxotrophic pressure, i.e., generating serine auxotrophs and coupling *serA* expression to one side of the toggle. This effect is illustrated for TOGGLE_green, with similar behavior obtained for TOGGLE_yellow (**Figure 3**). In the absence of auxotrophic pressure, the strains engineered with the original toggle switch (pECJ3) behaved as expected: IPTG addition triggered a complete switch to the mCherry state, which was stably maintained even after inducer withdrawal (**Figure 3A**). Only a small fraction of cells (∼1.5%) escaped and reverted to the GFP state, and it started escaping only after 15h without IPTG induction. By contrast, when *serA* dependence was introduced, the escape rate increased dramatically, with more than 90% of cells switching back to the GFP by the time we would expect the escape to start, based on the pECJ3 toggle behaviour (**Figure 3B**). Under these conditions, the toggle behaved as a unidirectional inducible system that naturally reverts once the inducer is withdrawn. This escape effect simplifies control by enabling unidirectional regulation with a single inducer.

**Figure 3.**
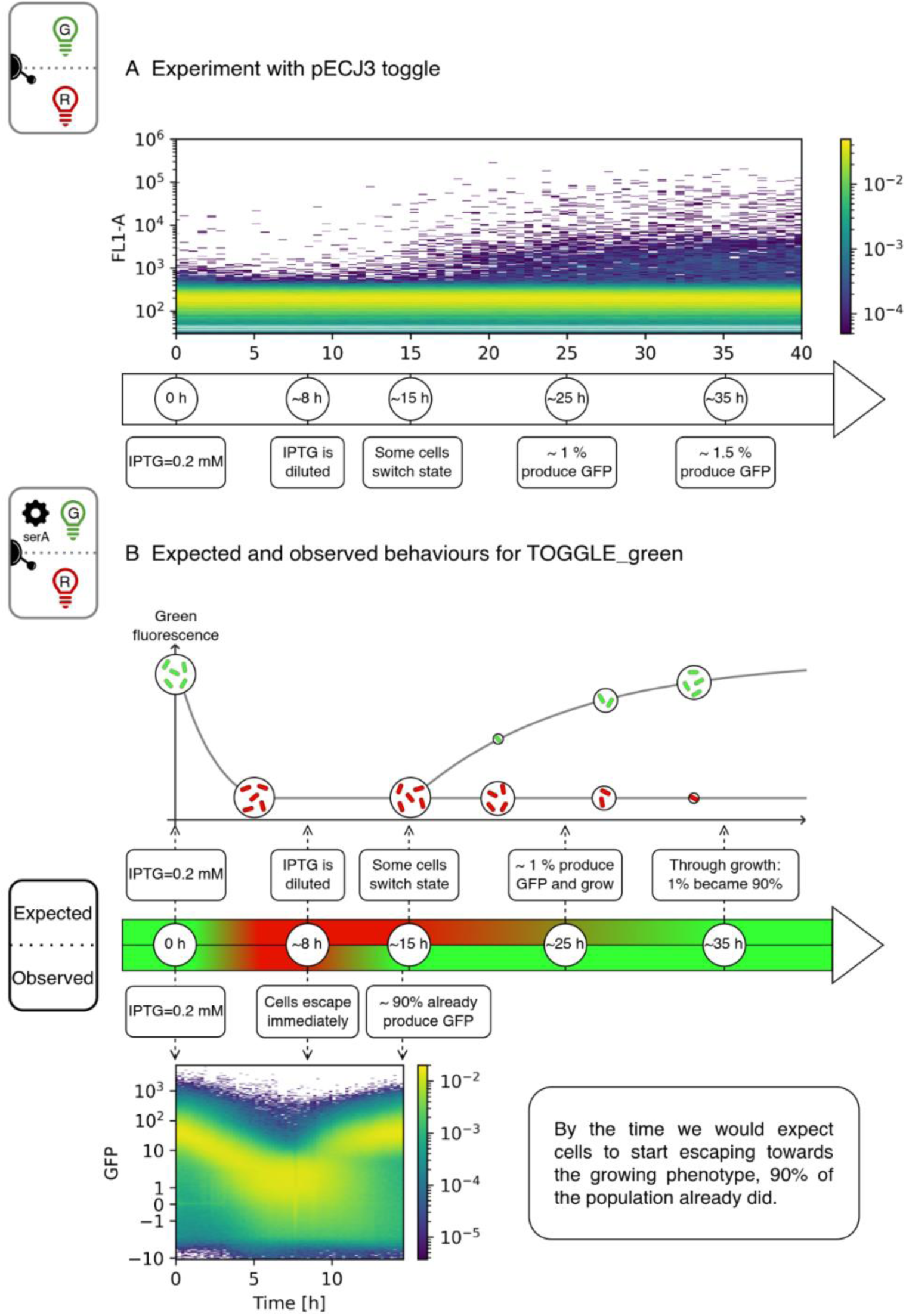
Auxotrophic pressure drastically alters population dynamics of the TOGGLE_green strain. **A** Automated flow cytometry (FC) profiling of the plasmid pECJ3 toggle after release from induction with 0.2 mM IPTG (FL1-A = GFP channel). Natural escape from the mCherry-producing phenotype occurs very slowly and stabilizes with ∼1.5% of the population producing GFP, while the remaining cells continue producing mCherry. **B** Based on the induction dynamics of pECJ3, when IPTG is used to induce the TOGGLE_green strain and inhibit SerA and GFP production, we would expect cells to remain trapped in the non-growing state for a long time. Instead, the SerA and GFP producing state reappears nearly immediately after IPTG is diluted.

### 3. Implementation of synthetic niches facilitate parameter inference for simulating mono-culture dynamics

As described above, introducing auxotrophic pressure caused the TOGGLE_green system to behave like a simple inducible gene circuit. To test this hypothesis, we cultivated the strain in a chemostat and applied IPTG pulses (**Figure 4A**). As expected, GFP levels decreased following IPTG addition, followed by a drop in biomass. However, once the inducer was diluted, the system naturally reverted to the opposite toggle state. This reversion was also reflected in biomass dynamics, with a natural increase in biomass after IPTG removal (**Figure 4B**).

**Figure 4.**
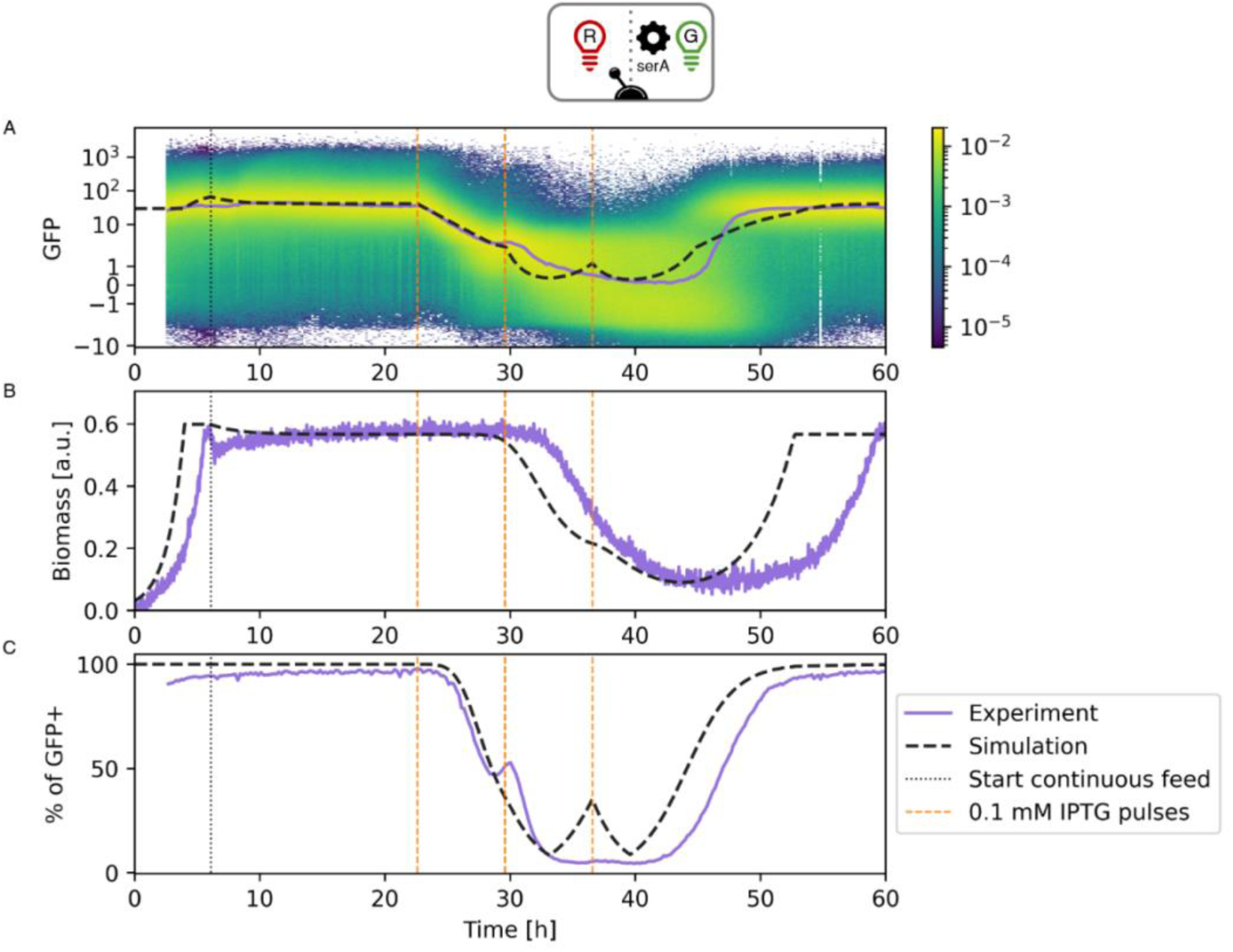
Auxotrophic pressure converts the TOGGLE_green circuit into a unidirectional inducible system. **A** Chemostat cultivation of TOGGLE_green with IPTG pulses showing GFP repression and natural reversion upon inducer washout. The experiment and simulation line represent the median fluorescence. **B** Biomass dynamics confirming growth recovery following IPTG removal. **C** Evolution of the percentage of GFP positive cells for the TOGGLE_green strain.

A simple ODE model incorporating a repression rule (see **Material and Methods**) successfully reproduced these dynamics, confirming that unidirectional control i.e., modulation with only one inducer, is sufficient to regulate co-culture composition (**Figure 4A–C**)

### 4. Increasing selective/auxotrophic pressure leads to mutational escape

To assess the robustness of the synthetic niche approach, we carried out continuous culture experiments at a dilution rate of 0.45 h⁻¹, testing each TOGGLE strain independently. Control was challenged by applying larger IPTG pulses (0.2 mM), triggered based on the fluorescence signal. In the TOGGLE_green strain, control efficiency deteriorated rapidly: after the second IPTG pulse, the system stopped responding as expected (**Figure 5**). This loss of responsiveness was evident both at the fluorescence level (**Figure 5A**) and in growth dynamics (**Figure 5B**). Subsequent IPTG pulses elicited progressively weaker *serA*–GFP responses, while the mCherry signal remained consistent with periodic induction. These experimental results strongly diverged from the simulations, highlighting a breakdown in the designed control logic. A similar decline in control efficiency was observed for the TOGGLE_yellow strain (**Supplementary Information**).

**Figure 5.**
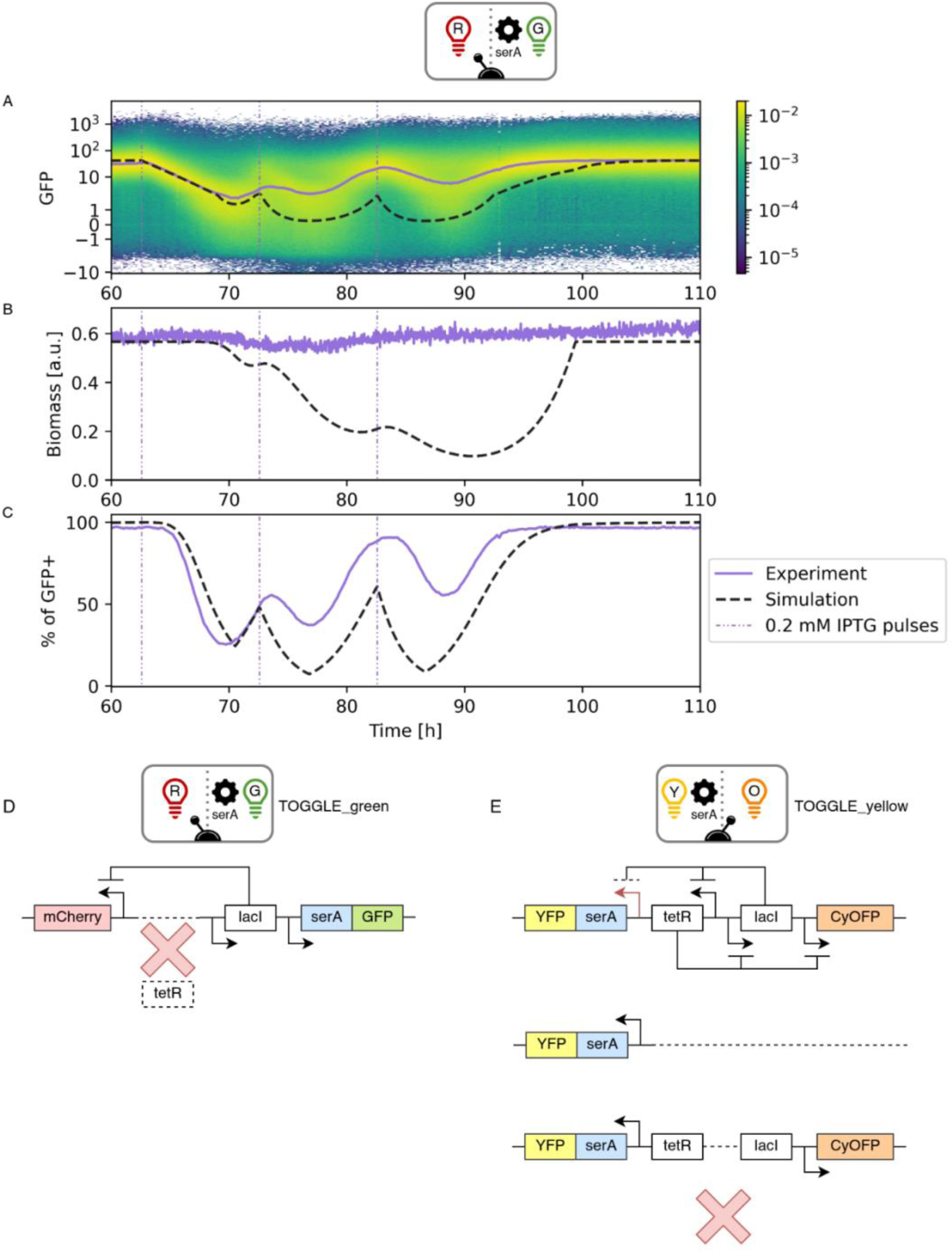
Mutational escape in TOGGLE strains under continuous culture. A–B. Breakdown of control efficiency in the TOGGLE_green strain after IPTG pulsing (0.2 mM, indicated by violet dashed lines). **A** Fluorescence dynamics determined based on automated FC showing rapid loss of GFP response while IPTG still induces mCherry (**Supplementary Figure S3**). The experiment and simulation lines represent the median of the fluorescence. **B** Corresponding growth curves highlight the decoupling of induction and growth. **C** Comparison of experimental fluorescence signals with model simulations, showing strong divergence after repeated pulses. **D** Sequencing of isolated colonies from TOGGLE_green revealed deletion of the *tetR* gene, consistent with the loss of GFP regulation and sustained mCherry expression. **E** The TOGGLE_yellow strain exhibited a similar decline in control efficiency but accumulated a broader spectrum of mutations, reflecting higher mutational diversity in this background.

To investigate the genetic basis of this failure, we isolated 5 colonies and performed plasmid extraction and sequencing. In the TOGGLE_green strain, no sample from the experiment described here was available for sequencing. However, sequencing of a subsequent experiment performed under strong IPTG induction revealed a deletion of the *tetR* gene (see **Supplementary Figure S5**, **Figure 5D**). Although this observation does not directly concern the experiment presented in this study, it indicates that genetic rearrangements can occur under similar induction conditions, and it fully explains the observed phenotype: loss of GFP response combined with IPTG-induced mCherry expression. Without *tetR*, the *serA*–GFP module is no longer repressed, effectively uncoupling regulatory control and allowing unchecked growth of the strain.

The TOGGLE_yellow strain exhibited a similar loss of control, but showed a markedly greater diversity of mutations, along with comparable diversity in fluorescence behaviours once these mutations had appeared. (**Figure 5E**).

### 5. Mutational escape impairs control of co-cultures

We next co-cultivated the TOGGLE_green and TOGGLE_yellow strains in a continuous bioreactor using the reactive flow cytometry protocol (Segregostat – **Figure 6**). As in monocultures, despite limiting the IPTG concentration, control efficiency deteriorated rapidly: after the first control cycle, the abundance of the TOGGLE_yellow strain dropped sharply. To test whether stronger induction could restore balance, we progressively increased IPTG pulse amplitudes, but the TOGGLE_yellow strain ratio continued to decline.

**Figure 6.**
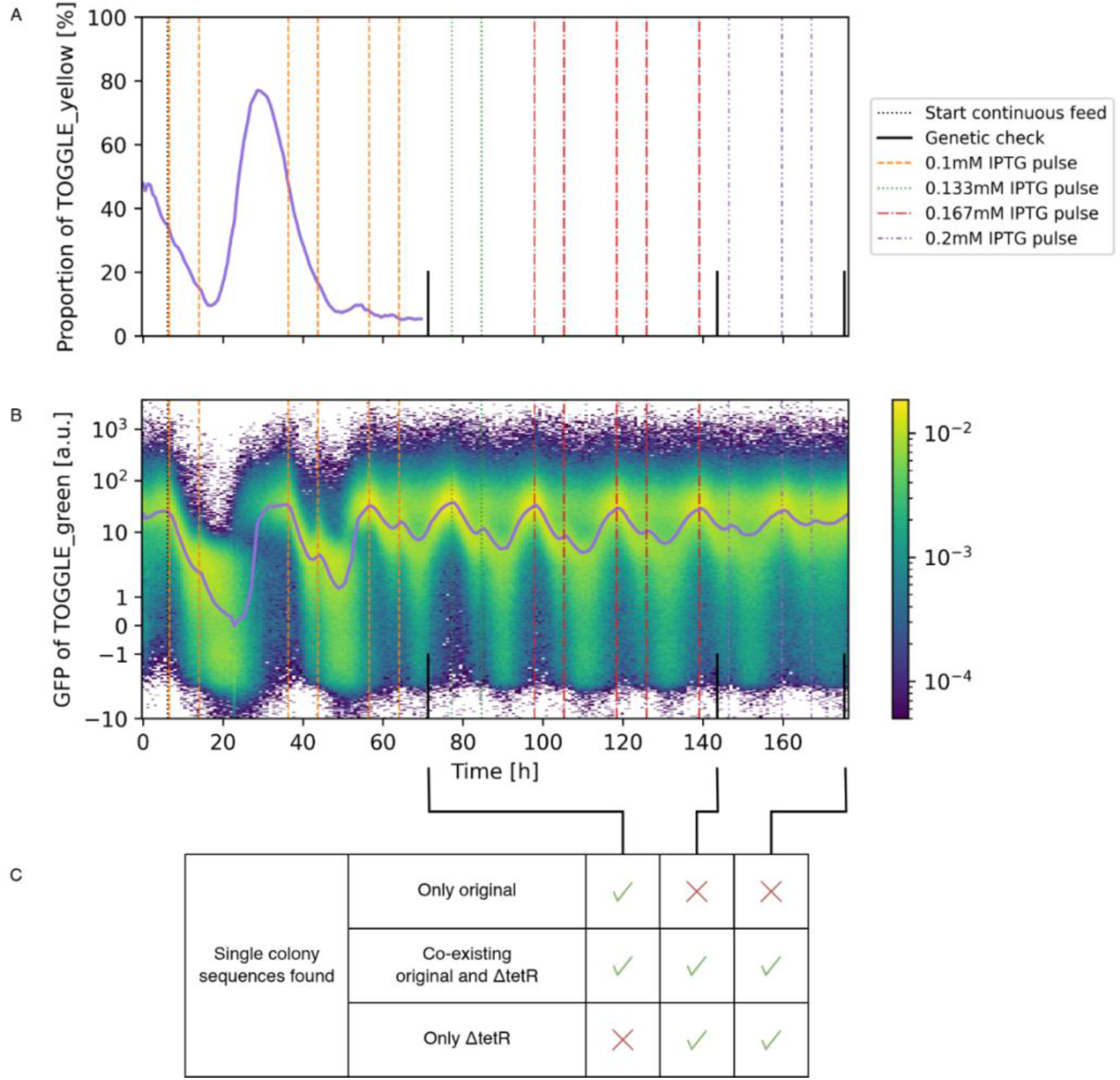
Mutational dynamics of TOGGLE strains in co-culture under selection pressure. A–B. Co-cultivation of TOGGLE_green and TOGGLE_yellow strains was performed in continuous bioreactors with Segregostat control. **A** Proportion of TOGGLE_yellow strain for as long as it can be meaningfully detected. Control is lost despite the low induction pressure, and the IPTG pulse concentration was increased to try and recover it. **B** GFP fluorescence of the TOGGLE_green population. GFP still responds to IPTG, but with a lower amplitude. **C** Sequencing and PCR analyses were carried out at different levels of selection pressure. Under low selection pressure, single-colony sequencing revealed a mixture of original TOGGLE_green cells, as well as cells carrying both wild-type and Δ*tetR* alleles. At high selection pressure and after extended cultivation, the population shifted toward coexistence of mixed alleles and complete Δ*tetR* mutants, with original non-mutated colonies largely eliminated. The schematic summarizes the distribution of colony types identified at each stage.

To uncover the genetic basis of this instability, we collected samples under low selection pressure at the start of cultivation and under high selection pressure at the end. Sequencing of several colonies in high-pressure samples revealed an approximately twofold reduction in read depth across the *tetR* region in all three reactors, suggesting partial loss of *tetR* sequences within single cells. PCR analyses confirmed this dynamic: under low selection pressure, both wild-type TOGGLE_green cells and mixed populations carrying mutated (Δ*tetR*) and non-mutated alleles were present. Under sustained high selection pressure, and after extended cultivation that removed most original TOGGLE_green cells, the population consisted mainly of cells harboring both wild-type and mutant sequences, along with a fraction of fully Δ*tetR* mutants.

## Discussion

This study establishes synthetic niches as a minimal and modular framework for regulating microbial co-cultures. By coupling cell growth to defined phenotypic states through toggle switch circuits, we effectively transformed metabolic compatibility into a controllable regulatory feature. Under moderate selective pressure, the system provided predictable and reversible control of strain abundance using a single inducer. The introduction of serine auxotrophy converted the bistable toggle into a unidirectional repression logic, simplifying the design while maintaining effective growth control. This architecture behaves functionally as a repressible rather than a bistable system, which may in fact facilitate deployment in industrial settings where simplicity and reliability are preferred [35,36].

These findings also suggest a design path in which the control circuit could be implemented in only one strain, the faster-growing or genetically tractable partner, while the other strain remains unmodified. Such asymmetric architectures would increase portability and reduce engineering effort, especially in cases where the slower or more specialized partner is difficult to transform. A notable example is yeast-bacteria co-cultures, where yeast is typically outcompeted by bacteria but has already been extensively engineered for bioproduction [13].

Compared with other approaches, such as cross-feeding, quorum-sensing, or toxin-antitoxin systems, synthetic niches impose minimal energetic burden and preserve metabolic independence of the production pathway, supporting the notion that population control can be achieved without extensive metabolic rewiring.

An additional strength of the synthetic niche framework lies in its quantitative transparency. Because the control dynamics can be described by a single repression rule, key parameters, such as induction thresholds, growth penalties, and recovery kinetics, can be measured on individual strains and then generalized to the co-culture context. This facilitates accurate model-based prediction of system behavior, in contrast with the empirical tuning required for ecological interactions or cross-feeding systems [10,14]. However, this predictability depends critically on the genetic integrity of the regulatory circuit: once mutations occur, parameter values lose meaning and the model rapidly diverges from experimental observations.

The principal limitation uncovered by this study is the evolutionary fragility of the control circuit. During extended continuous cultivation i.e., more than 80 generations at D = 0.45 h⁻¹, both toggle strains accumulated disabling mutations, most frequently deletions in the *tetR* region, leading to a complete loss of regulation. These mutations decoupled gene expression from growth, causing the escape of unregulated cells and collapse of co-culture control. Similar mutation-driven breakdowns have been reported in other burdensome expression systems [25,37], in which even minor growth disadvantages accelerate the fixation of escape variants. Our results show that this process can unfold within only a few control cycles under industrially relevant cell densities.

The probability of such genetic erosion scales with three parameters: the strength of selective pressure, which determines both the growth penalty and stress-induced mutation rate [38–43]; the population size, which sets the mutational target space [44,45]; and the number of generations, which defines total replication events [2,25]. Many published studies of synthetic consortia remain below these thresholds, operating with small populations or limited timescales that may mask instability. The continuous, high-density regime used here therefore provides a stringent benchmark for evaluating co-culture control methods and highlights the necessity of assessing genetic robustness under process-like conditions.

Several avenues could mitigate the instability observed here. Lowering the circuit burden, through tighter promoters, improved repression dynamics, or decoupling growth control from essential metabolic pathways, should reduce selection for escape. Alternative architectures could also be explored, including growth control via conditional toxin expression [17–19], optogenetic feedback [18], or hijacking of endogenous cell-cycle arrest modules [37]. Another promising approach would be to impose regulation only on the dominant strain, thereby relaxing selection across the population. Dynamic rejuvenation cycles or periodic strain replacement, already employed in long-term production systems [25,46], could further delay fixation of mutants. More broadly, integrating circuit design with evolutionary modeling and monitoring should allow the development of “evolutionarily informed” synthetic niches that sustain functionality over industrially relevant timescales.

Overall, our findings emphasize that evolutionary robustness must become a central design criterion in synthetic co-culture engineering. Synthetic niches offer a powerful and generalizable means to modulate strain ratios with minimal metabolic interference, yet their long-term success hinges on maintaining genetic stability. We advocate that validation of co-culture strategies should include extended continuous cultivations where both population size and generational span approach those of real bioprocesses. Only through such intensified and longitudinal testing can we reveal the true durability of control mechanisms. Combining these long-term assays with automated single-cell monitoring platforms such as the Segregostat [28,29] and real-time adaptive control [18] may ultimately yield microbial communities whose composition can be both programmed and evolutionarily sustained, enabling the transition from proof-of-concept consortia to industrially reliable bioprocesses.

## Material and methods

### 1. Genetic constructs

The pECJ3 plasmid was used in a *Escherichia coli* MG1655 Δ*lacAYZI* background [31]. All modified DNA sequences are provided with relevant primers in **Supplementary File genetic_constructs.7z**. To generate TOGGLE_green, the native *serA* coding sequence from MG1655 was cloned into the pECJ3 backbone directly downstream of the P_LtetO-1_ promoter and upstream of the ribosome binding site (RBS) used for GFP in the original construct (**Figure 2A, 2B**). The TOGGLE_yellow plasmid was designed following the same strategy, except that GFPmut3b (which we name GFP for convenience) and mCherry on pECJ3 were first replaced by CyOFP1 [47] (CyOFP) and sYFP2 [48] (YFP), respectively (**Figure 2D**). Both plasmids were inserted in an *Escherichia coli* MG1655 Δ*lacAYZI* Δ*serA* strain.

For TOGGLE_yellow, the native promoters controlling *serA* and YFP did not provide sufficient repression of growth. To reduce transcriptional leakage, the promoter was replaced by a P_LlacOsys_ promoter variant, obtained by merging the P_LlacO-1_ promoter [32] with the symmetric lacOsys operator sequence [33,34] (**Figure 7**).

**Figure 7.**
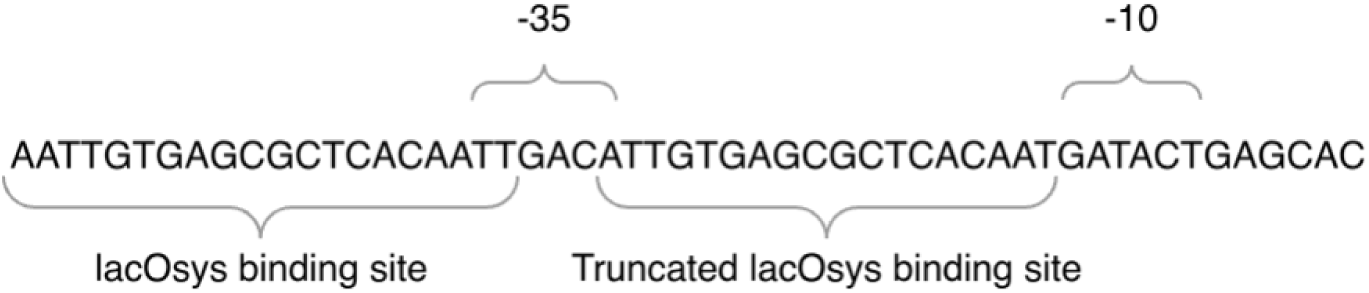
Sequence of the P_LlacOsys_ promoter.

### 2. Culture medium

Unless otherwise specified (e.g., inducer supplementation and concentration), the following medium was used to culture *E. coli* strains: a mineral medium containing (in g·L⁻¹), K_2_HPO_4_: 14.6, NaH_2_PO_4_⋅2H_2_O: 3.6, Na_2_SO_4_: 2, (NH_4_)_2_SO_4_: 2.47, NH_4_Cl: 0.5, (NH_4_)_2_-H-citrate: 1, glucose: 5, thiamine: 0.01. The medium was supplemented with 11 mL·L⁻¹ of a trace element solution. This solution was assembled from the following solutions (in g·L⁻¹): 3/11 of FeCl_3_⋅6H_2_O: 16.7, 3/11 of EDTA: 20.1, 2/11 of MgSO_4_: 120, and 3/11 of a metallic trace element solution. The metallic trace element solution contains (in g·L⁻¹): CaCl_2_⋅2H_2_O: 0.74, ZnSO_4_⋅7H_2_O: 0.18, MnSO_4_⋅H_2_O: 0.1, CuSO_4_⋅5H_2_O: 0.1, CoSO_4_⋅7H_2_O: 0.21. In bioreactors, the medium was supplemented with TEGO® Antifoam KS 911 (100 µL·L⁻¹). For plasmid maintenance, kanamycin sulfate (Gibco™, ≥ 95% purity) was added to the medium at a final concentration of 50 mg·L⁻¹. The trace element solution, thiamine, and kanamycin stock were filter-sterilized (0.22 µm) and added aseptically after autoclaving. All other medium components were heat-sterilized at 121 °C prior to supplementation.

### 3. Precultivation

Pre-cultures for both BioLector II and bioreactor experiments were prepared using the same procedure. Single colonies were inoculated into mineral medium containing 10 g·L⁻¹ glucose and cultured until OD > 5, after which glycerol stocks were prepared by mixing the culture 1:1 with sterile 50% glycerol. These glycerol stocks were then used to inoculate fresh mineral medium in baffled shake flasks, filled to 10% of their nominal volume, and incubated overnight at 37 °C at 150 rpm.

### 4. BioLector experiments

The BioLector II (m2p-labs, Germany) is a high-throughput cultivation platform enabling parallel cultivation of microbial strains, with online monitoring of growth and gene expression under well-aerated conditions that mimic small-scale stirred-tank bioreactors [49]. To characterize the growth of each phenotype independently, strains were first cultured under standard conditions for at least 9 h and then pre-induced overnight to force commitment to a single phenotype. Phenotype uniformity was verified before and after induction using an Attune NxT flow cytometer (Thermo Scientific™, blue and yellow lasers, 530/30 or 510/10 filters were used to detect the growing TOGGLE_green, and 530/30 or 540/20 were used to detect the growing TOGGLE_yellow).

For TOGGLE_green, induction was performed with 150 ng·mL⁻¹ anhydrotetracycline (aTc) (Thermo Scientific™, ≥ 98.0% purity), used when induction time had not yet been optimized, or with 1 mM IPTG (IPTG dioxane-free, Thermo Scientific™, ≥ 99.0% purity). For TOGGLE_yellow, induction was achieved using 25 ng·mL⁻¹ tetracycline hydrochloride (Tc) (Gibco™, ≥ 95% purity) or 1 mM IPTG. After induction, cultures were washed twice (centrifugation and resuspension) with glucose-free mineral medium to remove residual inducers, then inoculated into 48-well FlowerPlates (1 mL per well) to an initial OD600 of approximately 0.5 and a glucose concentration of 5 g·L⁻¹. For each strain, the plate layout included: 3 wells with medium only (baseline and negative control); 3 wells without inducer; 3 wells with inducer (to verify the absence of phenotypic escape). Because no difference in growth or fluorescence was observed between wells with and without inducer (Tc for the GFP- and YFP-expressing states, IPTG for the others), and flow cytometry confirmed the absence of phenotypic escape, only the non-induced conditions are presented. Biomass (scattered light) and fluorescence (520/25 filter, detecting both GFP and YFP signals) were recorded in a BioLector operated at 37 °C, 85% humidity, and 1200 rpm. Additional validation experiments are provided in the **Supplementary Information**.

### 5. Continuous bioreactor experiments

A detailed list of all continuous bioreactor experiments, including raw data and results, is provided in the **Supplementary Information**. The supplementary information also include additional experiments performed with the TOGGLE_yellow strain and complementary experiments confirming the basic behavior of the strains and co-cultures without induction. All bioreactor experiments were carried out at 37 °C and pH 7. Each experiment started with a batch phase until complete glucose depletion, followed by a continuous phase at a dilution rate of approximately 0.45 h⁻¹.

The experiment shown in **Figure 3A** was performed in a DASGIP system (original from the DASGIP group, bought in Europe, equivalent to the current DASbox system from Eppendorf, equipped with an OXY-4 mini system and OXYPro® oxygen probes from PreSens and EasyFerm Plus PHI K8 pH probes from Hamilton) using three vessels run in parallel (working volume 160 mL) (hence these are in triplicate). After an initial batch phase using 20 mL of pre-culture and no IPTG, the culture was shifted to a continuous phase for 17.25 h (>10 generations) with medium containing 0.2 mM IPTG, followed by a continuous phase of more than 75 h without IPTG. Forty hours of this IPTG-free phase are shown in **Figure 3A** for one representative replicate, as no further change occurred afterward. Online monitoring was performed via a custom cell-machine interface coupled to a BD Accuri C6 flow cytometer (GFP filter 510/15) [50][51]. Because the Accuri C6 is less sensitive than the Accuri C6+ and cannot accurately detect mCherry, the absolute GFP calibration used for other bioreactors could not be applied here, and raw FL1-A signals are reported.

All other monoculture experiments were performed in a 1 L Bionet F1 bioreactor (Bionet, Murcia, Spain, equipped with EasyFerm Bio PHI Arc pH sensors and VisiFerm RS485-ECS optical oxygen sensors from Hamilton). After an initial batch phase using 50 mL of pre-culture and no inducer, an overnight continuous phase without inducer was maintained to reach steady state. Pre-programmed inducer pulses were then applied (0.1 or 0.2 mM IPTG, 15 or 30 ng·mL⁻¹ Tc·HCl). Biomass was monitored in situ using a Dencytee Arc sensor (Hamilton). Glucose concentrations were determined by sampling, centrifugation, freezing of the supernatant, and analysis using the Megazyme D-Glucose HK assay kit. Flow cytometry monitoring was performed with a BD Accuri C6+ cytometer (4-blue laser configuration; filters 510/15, 540/20, 610/20, 565/20), except for the experiment shown in **Figure 3B**, which used filters 510/15, 540/20, 670LP. This alternative configuration accurately measured GFP and mCherry but could not discriminate mCherry from CyOFP1. All monoculture experiments in the Bionet system were performed in duplicate, except the experiment in **Figure 3B**, where pulse intervals were sufficiently long to avoid overlap, allowing three consecutive control sequences on the same culture, all yielding similar results. The conclusions from this experiment are consistent with those from other monoculture experiments but are particularly apparent here.

Co-culture experiments were carried out in an other DASGIP system (Eppendorf, bought in Belgium) using three vessels run in parallel (working volume 400 mL, all vessels were equipped with EasyFerm Plus PHI K8 pH probes from Hamilton) hence the co-culture experiments were run in triplicate. Flow cytometry measurements were collected using a BD Accuri C6+ cytometer (4-blue laser configuration; filters 510/15, 540/20, 610/20, 565/20). After inoculation with 20 mL of pre-culture, the batch phase was immediately followed by a continuous phase with strain-ratio control through IPTG pulsing (controls without induction are shown in the **supplementary information**). A pulse size of 0.1 mM IPTG was selected to minimize mutation. Pulses were triggered whenever the TOGGLE_yellow fraction dropped below 50%, with constraints of at most one pulse every 7 h and at most two pulses within 20 h. These constraints were chosen to allow sufficient dilution of IPTG to limit mutational escape and were validated through modeling.

### 6. Bioreactor data treatment

The code used for data processing is available at https://gitlab.uliege.be/mipi/published-software/2025-syntheticniches and uses Snakemake [52]. Flow cytometry data were analyzed using Accuri C6+ reference measurements for each individual fluorescent phenotype and for a non-fluorescent *E. coli* MG1655 strain. Based on these reference signals, each event was decomposed into autofluorescence, GFP, mCherry, YFP, and CyOFP contributions. Fluorescence values were then scaled so that 95% of the events from the non-fluorescent MG1655 reference strain fell below a value of 2 for each fluorophore channel. The value of 2 was therefore used as the detection threshold relative to the wild type autofluorescence background. Note that the expression of one fluorophore slightly increases the baseline signal detected in other channels (**Figure 8**). Negative values may appear after autofluorescence subtraction.

**Figure 8.**
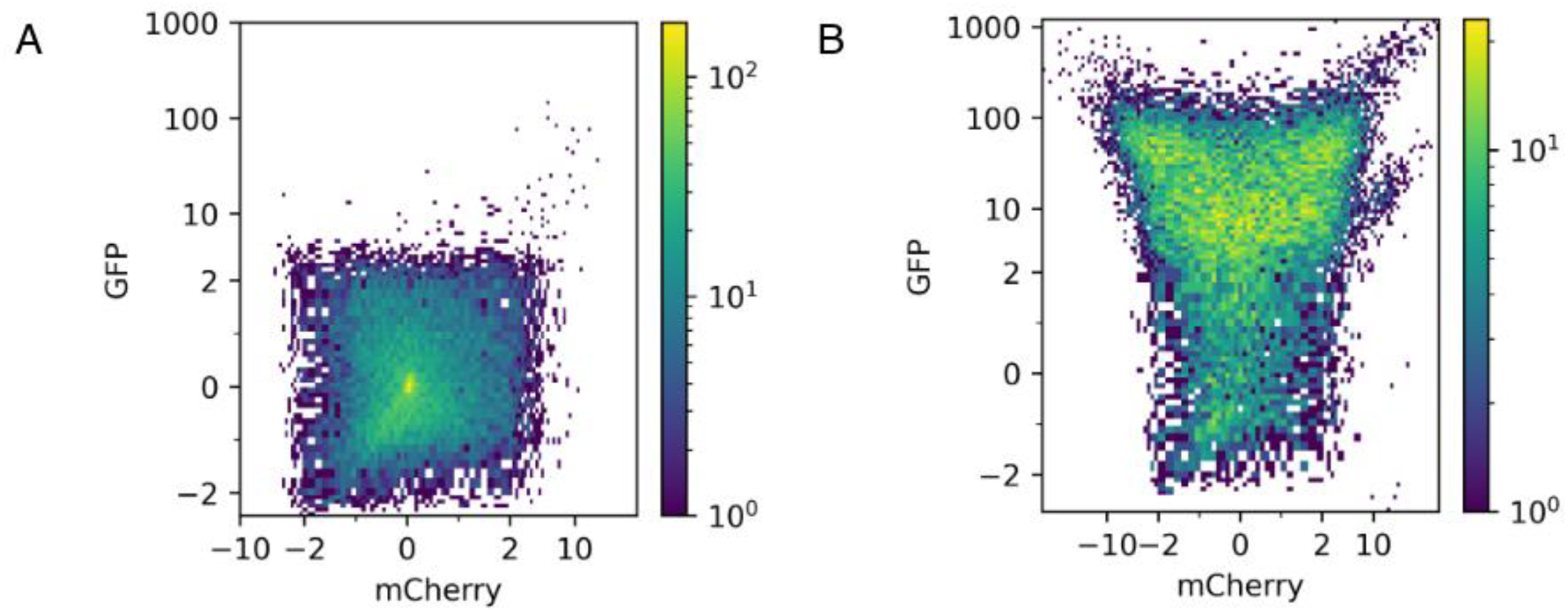
Transformation of a flow cytometry sample of A MG1655 (wild type) and B TOGGLE_green expressing GFP. Light yellow areas indicate a high number of cells, while dark blue area indicate low cell density. The [-2,2] interval is where 95% of the cells are present; 2 was an arbitrarily defined threshold. For the few percent of cells with low GFP fluorescence for TOGGLE_green, the mCherry fluorescence is also within [-2,2]. However, for high GFP values, noise produces cells located in a log fluorescence range between 2 and -2 in the mCherry channel. Therefore, a diagonal line is used to separate phenotypes instead of a threshold.

Distinguishing phenotypes in the TOGGLE_green strain was straightforward. Upon IPTG induction, GFP decreases first, followed by an increase in mCherry. A TOGGLE_green cell was therefore classified as GFP-positive (and SerA-positive) when GFP > 2 and GFP > mCherry. For the TOGGLE_yellow strain, CyOFP increases before YFP reaches its basal level, presumably because CyOFP is detected more easily than mCherry with a 488 nm excitation. Since only low SerA expression is required for growth (consistent with the need for a very tight promoter), a threshold was defined to isolate a pure CyOFP phenotype; any cell above that threshold was considered growth competent. Practically, a TOGGLE_yellow cell was defined as YFP-positive when YFP > 0.38 × CyOFP and YFP > 1.

In co-cultures, strains were distinguished based on the fluorophore vector space: cells closer to the autofluorescence–GFP–mCherry plane were assigned to TOGGLE_green, and cells closer to the autofluorescence–YFP–CyOFP plane to TOGGLE_yellow. Because CyOFP and mCherry emit weakly under 488 nm excitation, monoculture controls confirmed partial overlap between CyOFP and mCherry signals. This overlap was expected, as these fluorophores correspond to phenotypes that should not coexist during co-culture: under IPTG, mCherry (induced), GFP (repressed but growing), and YFP (induced and growing) are present; under Tc, CyOFP (induced), YFP (repressed but growing), and GFP (induced and growing) are present. Both inducers are never applied simultaneously, as this would activate both sides of the toggle switch. Consequently, in IPTG-controlled co-cultures, all cells that could correspond to either CyOFP or mCherry were classified as mCherry. Verification using monocultures confirmed that this classification identified more than 90% of events correctly in all cases, and more than 95% except under prolonged or strong induction.

All monoculture and co-culture experiments were performed on flow cytometers of the same model. A calibration was therefore applied to convert co-culture measurements into their equivalent monoculture measurement space (correlation of the shape y=a·x^b^+c between the intensity of identical samples passed in both cytometers, for all used channels, with the various fluorescent proteins, individual correlation points were found by comparing identical measurement percentiles between machines).

### 7. Model parameter estimation

A complete description of the parameter inference procedure is provided in the **Supplementary Information**. Briefly, growth of the productive (growing) phenotype was described using a classical Monod equation:

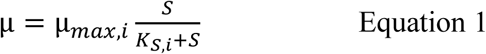

where μ is the instantaneous growth rate a strain, μ_max, i_ is its maximum growth rate, 𝑆 is the glucose concentration, and 𝐾_𝑆,𝑖_ is the glucose concentration at half-maximum growth.

For the non-growing phenotype, we considered:

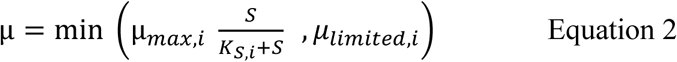

where μ_𝑙𝑖𝑚𝑖𝑡𝑒𝑑,𝑖_ represents the maximum growth rate of the non-growing phenotype of strain i. Maximum growth rates were not identical in Biolector and bioreactor batch. Therefore, we deduced the growth rates (μ_𝑚𝑎𝑥,𝑖_, μ_𝑙𝑖𝑚𝑖𝑡𝑒𝑑,𝑖_) of individual from the biomass evolution during a series of 0.1 mM pulses for TOGGLE_green and during a series of 15 ng/mL Tc pulses for TOGGLE_yellow, while taking into account the measured phenotype ratio over time. 𝐾_𝑆,𝑖_ was determined using the residual glucose concentrations at steady-state. The biomass yield was determined based on the biomass obtained in batch and the provided glucose concentration. Besides the maximum growth rates, growth parameters were considered equal between phenotypes and different between strains.

The induction parameters were determined by analysing the fluorescence and phenotype ratio behaviour based on the inducer concentration, and assuming the previously determined growth parameters are correct. For TOGGLE_green, the GFP median of the population could be reproduced by an IPTG-repressible production equation:

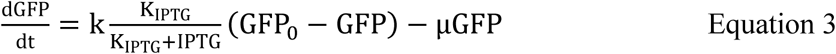

where GFP is the GFP concentration, 𝐾_IPTG_is the repression constant, IPTG is the IPTG concentration in the reactor, 𝑘 is the production constant, GFP_0_ is the GFP level at which production stops, μ is the growth rate.

### 8. Detection of mutational escape

To detect and isolate mutations, culture samples were withdrawn from the bioreactor and immediately mixed 1:1 with sterile 50% glycerol before storage at −80 °C. Frozen samples were subsequently streaked on LB agar, and single colonies were picked for liquid culture. Plasmid DNA was extracted using the corresponding commercial miniprep kit (NucleoSpin Plasmid EasyPure by Macherey-Nagel) and sequenced by Eurofins Genomics (whole plasmid sequencing).

To determine the presence or absence of the *tetR* gene, PCR was performed on colonies obtained from the LB plates (six colonies per time point and per biological replicate). One PCR used a primer located inside *tetR*; no amplification indicated loss of the gene (primers: GGCGAGTTTACGGGTTGTTA and GGCATACTCTGCGACATCGT). A second PCR amplified the genomic region encompassing *tetR*, yielding a long amplicon when *tetR* was present and a shorter fragment when *tetR* was absent (primers: GGCATACTCTGCGACATCGT and TTGGTCACCTTCAGCTTGG). Colonies containing both the wild-type and the deleted allele produced two amplicons.

### 9. Data availability

All code used for this paper is available at https://gitlab.uliege.be/mipi/published-software/2025-syntheticniches, and all data is available at https://doi.org/10.5281/zenodo.17663997.

## Acknowledgements

We would like to thank Fabian Moreno-Avitia for helping to create the concept of this project. We would also like to thank Maximilian Tieke for providing the basis of **Figure 1C**. VV and LH were supported by a FRIA grant provided by the “Fonds de la Recherche Scientifique” FRS-FNRS from the Walloon region of Belgium. The author are also very grateful to the Feder program 2021–2027, project PHENIX FoodBooster ULiege-SPW for the acquisition of an extended version of the Segregostat platform.

## Author contributions

FD and JAM supervised the study. VV, FD, JAM and LH contributed to the conceptualization and methodology of the investigation, and the experiments were performed by VV, ST, AZ and LH. Data analysis was performed by VV, with parts of the software written by JAM. FD, LJ and VV wrote the manuscript

## Declaration of interests

The authors declare no competing interests.

## Notes

### Competing Interest Statement

The authors have declared no competing interest.

