## Supplementary material for "Synthetic niches enable co-culture bioprocessing but are prone to mutational escape"

First version

Vincent Vandenbroucke, Juan Andrés Martínez, Lucas Henrion, Andrew Zicler, Samuel Telek, Laurie Josselin, and Frank Delvigne[1]

Terra Research and Teaching Centre, Microbial Processes and Interactions (MiPI), Gembloux Agro-Bio Tech, University of Liège, Gembloux, Belgium

24th November 2025

**3 Supplementary file list** **17**

### 1 Experiments performed

TABLE 1 below reports the list of the experiments carried out in the context of this article and their main results. It is followed by additional practical information about some of these experiments where necessary, and results mentioned in the manuscript concerning TOGGLE_yellow.

Table 1: List of experiments performed

| | Experiment | Figure | Strain and Culture device | Induction profile | Main results | Additional note | Relevant section in main text |
|---|---|---|---|---|---|---|---|
| 1 | 2022-09-26-serineSweep | | Δ*serA E.coli* in Biolector | Culture with varying amounts of serine supplementation. | The final biomass is directly proportional to amount of serine provided. | Note 1 | 1. Synthetic niches enable the selective control of the growth rate of microbial species involved in co-cultures |
| 2 | 2023-07-22-inductionSweep | | pECJ3 toggle in Biolector | Pre-culture without induction and culture with range of IPTG and Tc concentration. | Measured induction range for IPTG and Tc and verified no growth reduction with Tc at induction concentrations. | | |
| 3 | 2023-07-09-TSGCharacterisation | 2A | TOGGLE_green in Biolector | Induction with one inducer in pre-culture, no inducer in culture. | Measured growth rates of isolated phenotypes. | | |
| 4 | 2024-02-22-TSY2Characterisation | 2B | TOGGLE_yellow in Biolector | | | | |
| 5 | 2023-07-23-TSGCross-feeding | S1A | TOGGLE_green in DasBox equivalent (triplicate) | Continuous culture with inducer towards growth. Once at steady-state, the culture was supplemented with Δ*serA E.coli* and the resulting strain ratio was monitored. | The TOGGLE_green strain did not meaningfully export nutrients for another strain to survive. | Note 2 | |
| 6 | 2024-07-25-TSY2Cross-feeding | S1B | TOGGLE_yellow in DasBox equivalent (triplicate) | | The TOGGLE_yellow strain enabled the growth of ~1-2% of Δ*serA E.coli.* | | |
| 7 | 2024-10-16-TSBaselineNoise | 3A | pECJ3 toggle in DasBox equivalent (triplicate) | Batch without induction, continuous culture overnight with induction 0.1 mM IPTG, then no more induction. | After the release of induction, at most 1.5% of the population changed state through noise. | | 2. Auxotrophic pressure induces cell escape from the toggle switch |
| 8 | 2023-10-26-TSGPulseTest | 3B, S2 | TOGGLE_green in Bionet | Series of 3 0.2 mM IPTG pulses, then 3 0.1 mM pulses, spaced wide enough that the culture came back to its steady-state between pulses. | Individual pulses over time had reproducible effects, and there is no need for the second inducer. | Note 3 | |
| 9 | 2024-08-08-TSGPulses1 | | TOGGLE_green in Bionet | Series of 0.1 mM IPTG pulses, then series of 0.2 mM pulses. | Measured induction and growth parameters in continuous culture, and observed reduced effectiveness of biomass control during the second pulse series. For TOGGLE_yellow, mutations were checked and detected at the end of the culture. | Note 4 | 3. Implementation of synthetic niches facilitate parameter inference for simulating mono-culture dynamics<br>And<br>4. Increasing selective/auxotrophic pressure leads to mutational escape |
| 10 | 2024-09-05-TSGPulses2 | 4, 5, S3 | | | | | |
| 11 | 2024-07-11-TSY2Pulses1 | S4 | TOGGLE_yellow in Bionet | Series of 15 ng·mL$^{-1}$ pulses, then series of 30 ng·mL$^{-1}$ pulses. | | Note 5 | |
| 12 | 2024-10-02-TSY2Pulses2 | | | | | | |
| 13 | 2025-01-30-TSGMutation | S5 | TOGGLE_green in Bionet | Continuous feed with i) no IPTG, ii) 0.1 mM IPTG, iii) no IPTG, and iv) 0.1 mM IPTG. | After the first loss and recovery of biomass, the Δ*tetR* mutation was detected, and the GFP was no longer regulated. | Note 6 | 4. Increasing selective/auxotrophic pressure leads to mutational escape |
| 14 | 2025-03-12-freeCoCulture | S6 | Co-culture in DasGIP (triplicate) | No inducer. | TOGGLE_yellow was, as expected based on growth rates, always the losing strain. | Note 7 | 5. Mutational escape impairs control of co-cultures |
| 15 | 2025-05-21-ControlledCoCulture | 6 | Co-culture in DasGIP (triplicate) | Pulses of inducer based on control rule. After TOGGLE_yellow was completely lost with 0.1 mM pulses, pulse concentration was gradually increased. | Despite reasonably low induction concentration, the TOGGLE_green mutated and escaped control. | | |

The following section contrains additional practical information about the experiments from Table 1 where necessary, and results mentioned in the manuscript concerning Toggle_yellow:

**Note 1, Biolector experiments:** The first biolector experiments were used to ensure the strains behaved as expected. In particular, the induction range of IPTG and tetracycline (Tc) were determined with the 2023-07-22-inductionSweep experiment.

**Note 2, 2023-07-23-TSGCross-feeding, 2024-07-25-TSY2Cross-feeding:** One potential concern when there are phenotypes that cannot produce an amino acid, is cross-feeding from producing cells towards non-producing. Serine is the basis of all amino acid metabolism in *E. coli*. Therefore, if the non-producing cells can scavenge serine, other amino acids, or cell debris produced by the growing phenotype in the culture, it could allow the non-growing phenotype to grow as well. To ensure that no cross-feeding effectively happened, growing cultures of Toggle_green and Toggle_yellow were supplemented with a Δ*serA* *E. coli* without plasmid. If there is any cross-feeding, then the Δ*serA* strain should be able to survive in the culture. To get a worst case scenario, the continuous culture was supplemented with 25 $ng \cdot mL^{-1}$ Tc for Toggle_green or 0.1 mM IPTG for Toggle_yellow. Pre-cultures in minimal medium with 1 $g \cdot L^{-1}$ serine (an excess of serine) were added to reach about 5% of the total events, enough to be measurable and credibly survive for long enough in case of cross-feeding, but not enough to meaningfully alter bioreactor concentrations in serine or other metabolites. For Toggle_green, no Δ*serA* survived, while for Toggle_yellow, 1 to 2% of Δ*serA* were sustained, indicating that, under normal conditions, the non-growing phenotypes should be nearly totally eliminated from the culture (Figure S1).

**Note 3, 2023-10-26-TSGPulseTest:** For 0.2 mM pulses, there was a small effect on biomass for each pulse and the fluorescence took longer than the biomass to come back to normal. Pulses were spaced 14.5 h apart. For 0.1 mM pulses, a single pulse was not enough to affect biomass. Pulses were spaced 12.5 h apart. A summary of the experiment is presented in Figure S2.

**Note 4, 2024-08-08-TSGPulses1, 2024-09-05-TSGPulses2:** As mentioned in the main text, even when control is lost over GFP production, mCherry still responds to IPTG. This was observed in all experiments with Toggle_green and a loss of control (Figure S3).

**Note 5, 2024-07-11-TSY2Pulses1, 2024-10-02-TSY2Pulses2:** A behaviour similar to Toggle_green was observed when pulsing Tc for Toggle_yellow. However, the concentrations of Tc employed were lower in the induction curve so that only a fraction of Toggle_yellow switched to the non-growing phenotype. When the induction pressure was high enough for all cells to switch, all cells switched to the non-growing phenotype, quickly switched back to growing, and stopped responding to tetracycline. Towards the end of the culture, one culture exhibited a population with constitutive YFP and CyOFP expression (depicted in Figure S4, while the other exhibited a small non-fluorescent population, which prompted the sequencing and revealed diverse mutations.

**Note 6, 2025-01-30-TSGMutation:** Based on the results with Toggle_yellow, we wanted to check for mutations with Toggle_green. An experiment with constant 0.1 mM induction (IPTG in the feed) showed initial wash-out, then recovery of the population (Figure S5). The resulting cell population still responded to IPTG by modulating mCherry production, but not by modulating

GFP, just like the reduced sensitivity in the previous repeated pulse experiments with the strain. Sequencing revealed the *tetR* deletion.

**Note 7, 2025-03-12-freeCoCulture:** A basic co-culture without induction was first performed, to verify that the expected strain was winning. As shown in FIGURE S6, TOGGLE_yellow is outcompeted in the co-culture, as expected due to their lower growth rate.

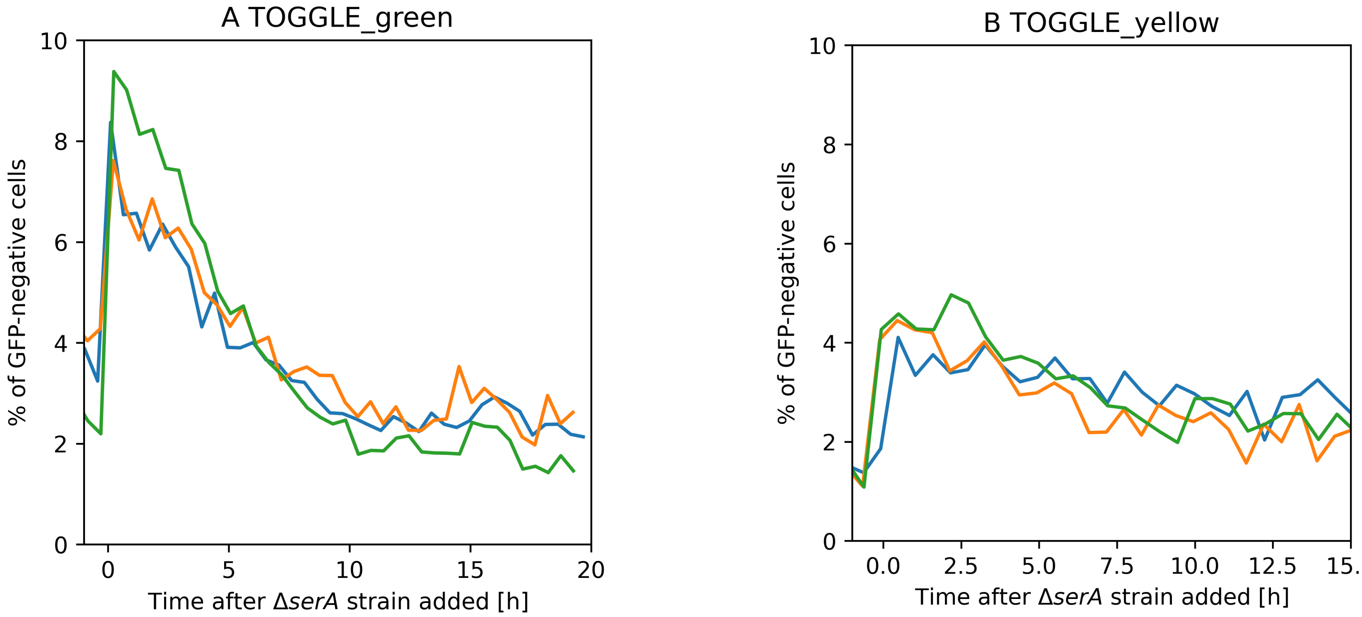

Figure S1: Evolution of GFP-negative population after injecting $\Delta$*serA E. coli* in a TOGGLE_green (**A**) or TOGGLE_yellow (**B**) culture to test for cross-feeding, in triplicate (experiments 5 and 6 in TABLE 1). **A** The measurement are reduced to background noise indicating that $\Delta$*serA E. coli* totally disappears from the culture. **B** The measurements get to a level slightly higher than background noise, indicating small amounts of cross-feeding.

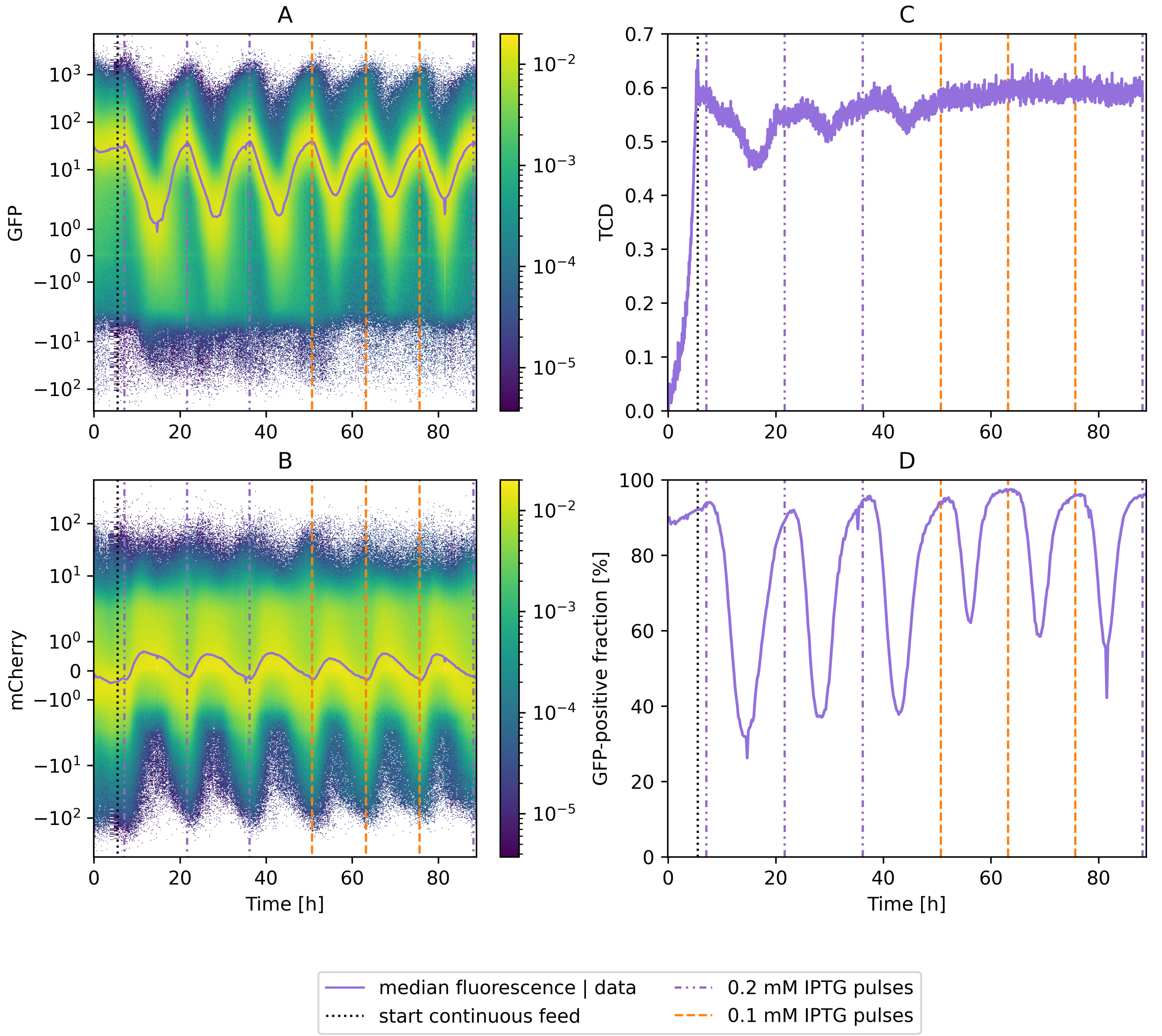

Figure S2: Summary of the 2023-10-26-TSGPulseTest experiment (row 8 in Table 1), referenced in paper section 2. **A** Evolution of GFP over time, the line indicates the median fluorescence. Light yellow indicates a high proportion of cells at a given time and fluorescence, dark blue indicates a low proportion. **B** Similarly, evolution of mCherry over time. **C** Measured cell density over time (from Hamilton Dencytee probes, 0.5 TCD≃5 OD). **D** Percentage of GFP-positive (growing) cells over time.

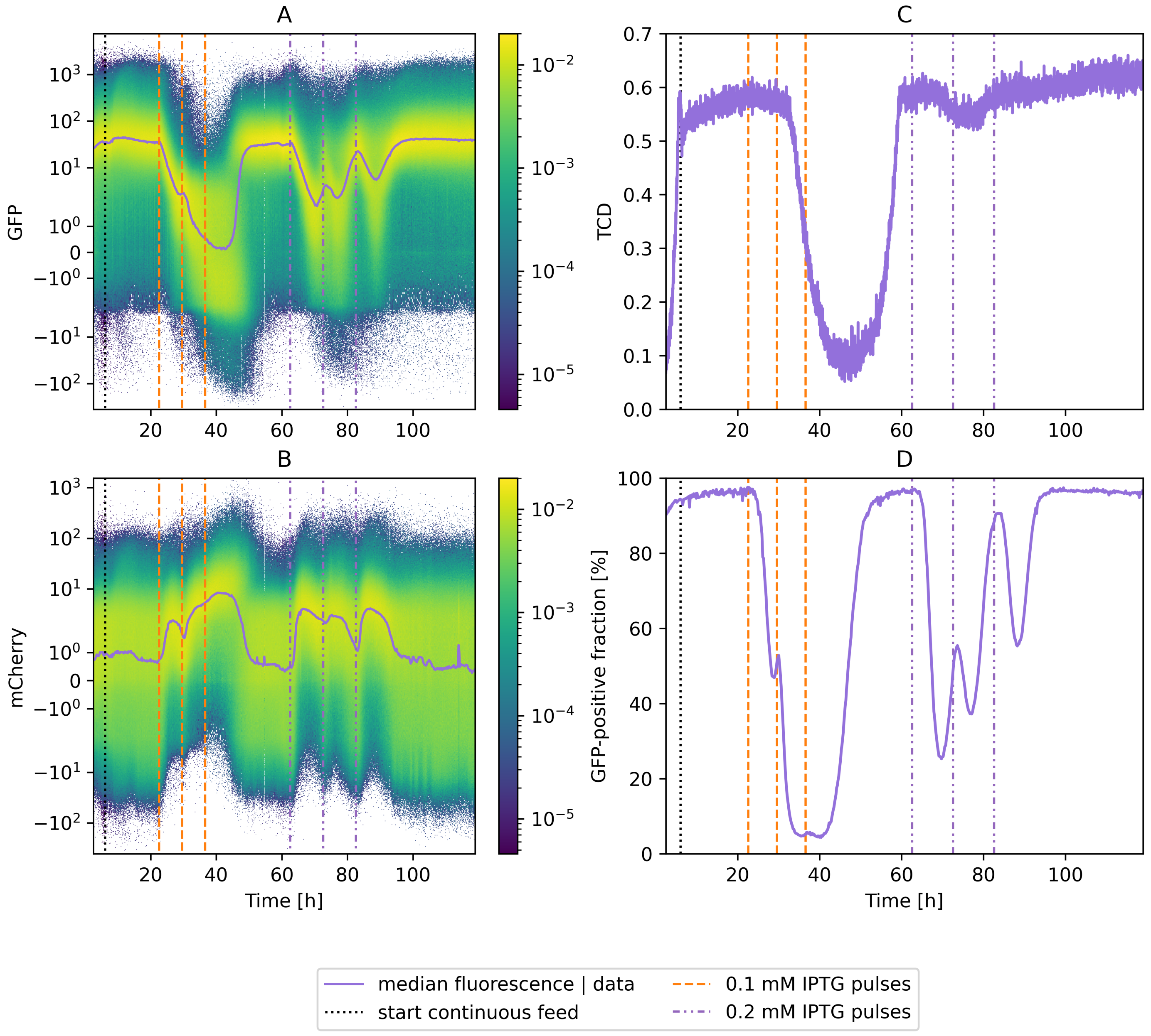

Figure S3: Summary of the 2024-09-05-TSGPulses2 experiment (row 10 in Table 1). Figure referenced in paper section 4. **A** Evolution of GFP over time, the line indicates the median fluorescence. Light yellow indicates a high proportion of cells at a given time and fluorescence, dark blue indicates a low proportion. **B** Similarly, evolution of mCherry over time. **C** Measured cell density over time (from Hamilton Dencytee probes, 0.5 TCD≃5 OD). **D** Percentage of GFP-positive (growing) cells over time.

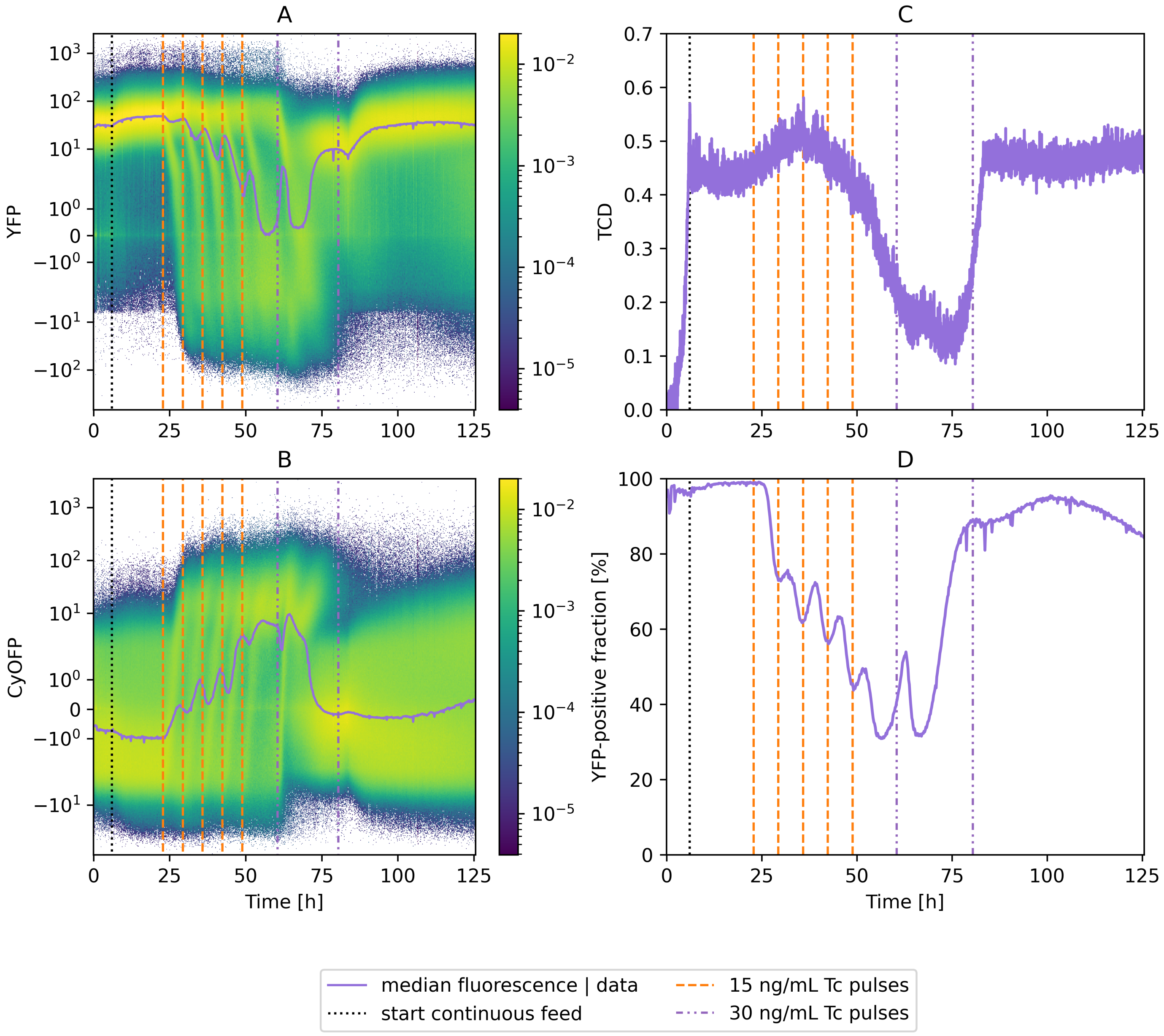

Figure S4: Summary of the 2024-07-11-TSY2Pulses1 experiment (row 11 in TABLE 1), mentioned in paper section 4. **A** Evolution of YFP over time, the line indicates the median fluorescence. Light yellow indicates a high proportion of cells at a given time and fluorescence, dark blue indicates a low proportion. **B** Similarly, evolution of CyOFP over time. Unlike TOGGLE_green, TOGGLE_yellow stopped responding to high concentrations of inducers, including for CyOFP. After the second pulse of 30 ng·mL$^{-1}$ Tc, a population exhibiting both and fluorescent proteins appeared in this experiment, causing a gradual increase in the CyOFP values. **C** Measured cell density over time (from Hamilton Dencytee probes, 0.5 TCD≃5 OD). **D** Percentage of YFP-positive (growing) cells over time.

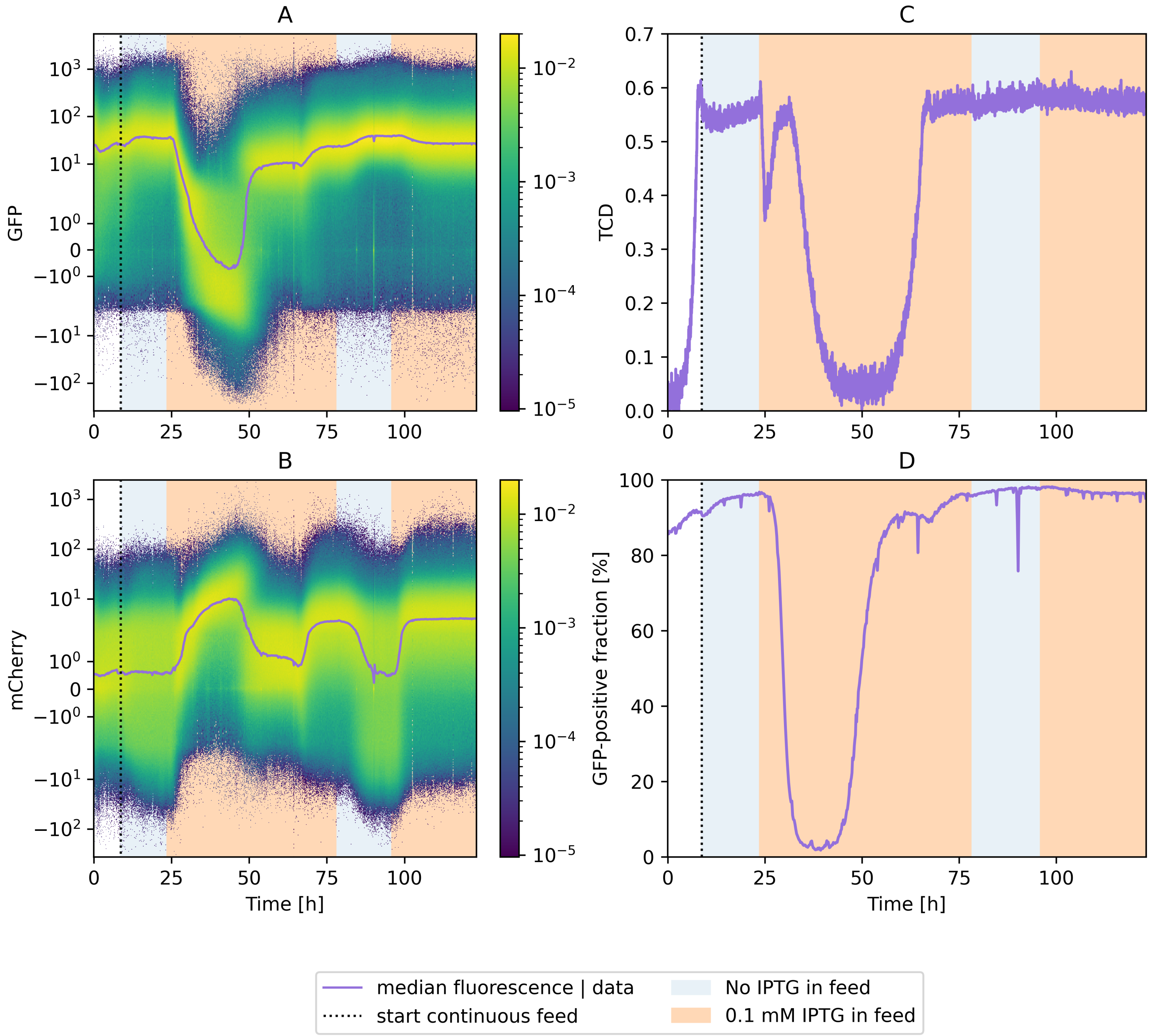

Figure S5: Summary of the 2025-01-30-TSGMutation experiment (row 13 in Table 1), mentioned in paper section 4. After a loss of biomass until 50 h of experiments, cells recover both growth capability and GFP expression, and a second indution period shows no loss of growth capacity. **A** Evolution of GFP over time, the line indicates the median fluorescence. Light yellow indicates a high proportion of cells at a given time and fluorescence, dark blue indicates a low proportion. **B** Similarly, evolution of mCherry over time. **C** Measured cell density over time (from Hamilton Dencytee probes, 0.5 TCD≃5 OD). Due to a technical error, the first hour with IPTG did not receive any carbon, hence the initial loss of biomass. **D** Percentage of GFP-positive (growing) cells over time.

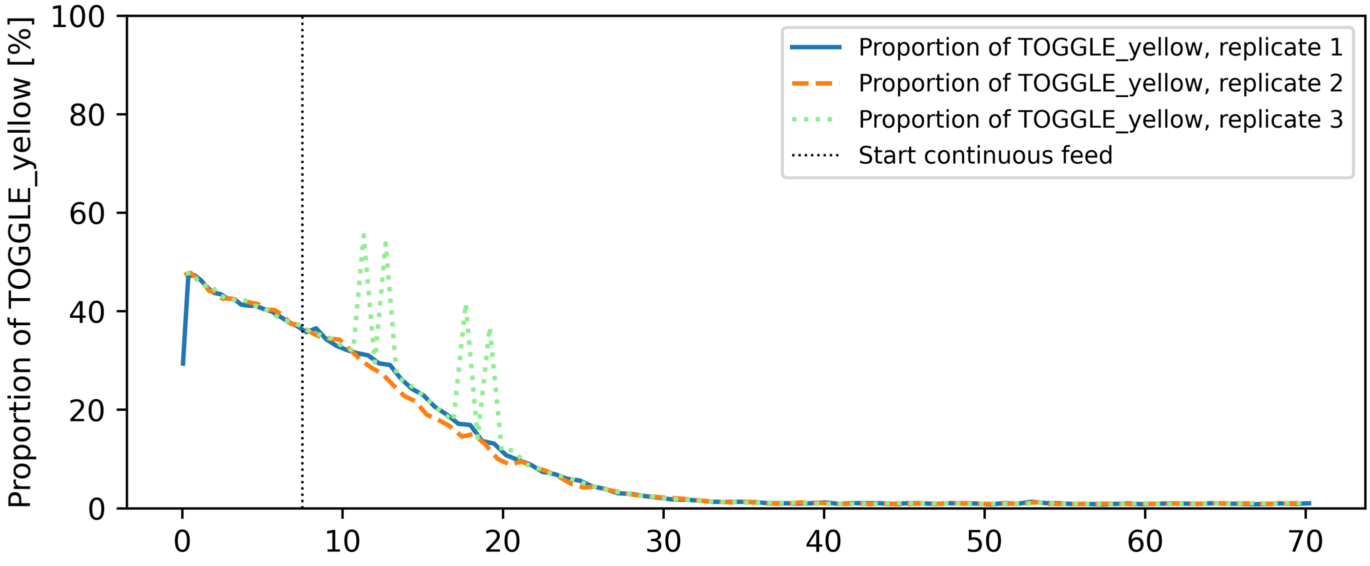

Figure S6: Proportion of Toggle_yellow in three co-cultures without inducer (2025-03-12-freeCoCulture, experiment 14 in Table 1).

### 2 Parameter inference to model the strains

#### 2.1 Modular modelling framework

When modelling in silico parameters, one common hurdle is the determination of multiple parameters with different units and meanings based on diverse sources of information. To infer these parameters, it is possible to define a single model of the system, define a loss function that summarises the various sources of information into a single error number, and minimise that loss function to find the best possible parameters for the system.

To create such a loss function, choices are necessarily made regarding the relative importance of different, unrelated parameters. For example, for the case of this paper, a model would include both fluorescence and biomass data, and a single loss function would have to weigh the error for these parameters against each other, leading to 'optimal parameters' depending on the loss function.

However, in practice, different measurements are associated to different parts of the process, and to different model parameters. Therefore, another approach is to consider the modelled system as a collection of smaller, independent models, each representing part of the process and accounted for independently from each other. This was the approach chosen here, as depicted in FIGURE S7: first, the growth parameters were estimated considering the fluorescence data as ground truth and optimising the model to fit biomass observations. Then, the growth model was used to determine a fluorescence model suitably representing reality. Finally, the full model was applied both to confirm it could represent all performed experiments – it was only valid when mutations were not present – and provide an estimation for the behaviour of a co-culture, with different inducer pulse profiles.

To achieve this modularity, the models were implemented using a custom interface (available at `https://gitlab.uliege.be/mipi/published-software/2025-syntheticniches`), allowing to define parts of the model separately and combine them as needed. This lets us easily replace the module containing the fluorescence of an experiment by a simulation of the fluorescence, or a pulse profile from an experiment by a control rule that uses the simulated fluorescence profile to determine when to add inducer. Differential equations were simulated using an Euler step of 0.01 h, which was sufficient to ensure numerical stability.

#### 2.2 Determination of growth parameters

For both TOGGLE_green and TOGGLE_yellow, the growth model considered was a classical Monod equation for the growing phenotype:

$$\mu_i = \frac{\mu_{\max,i} S}{K_{S,i} + S} \,, \tag{SE 1}$$

where $\mu_i$ is the normal growth rate for the strain $i$, $\mu_{\max,i}$ is the corresponding maximum growth rate, $S$ is the substrate (glucose) concentration in the bioreactor, and $K_{S,i}$ is the substrate concentration at half-maximum growth.

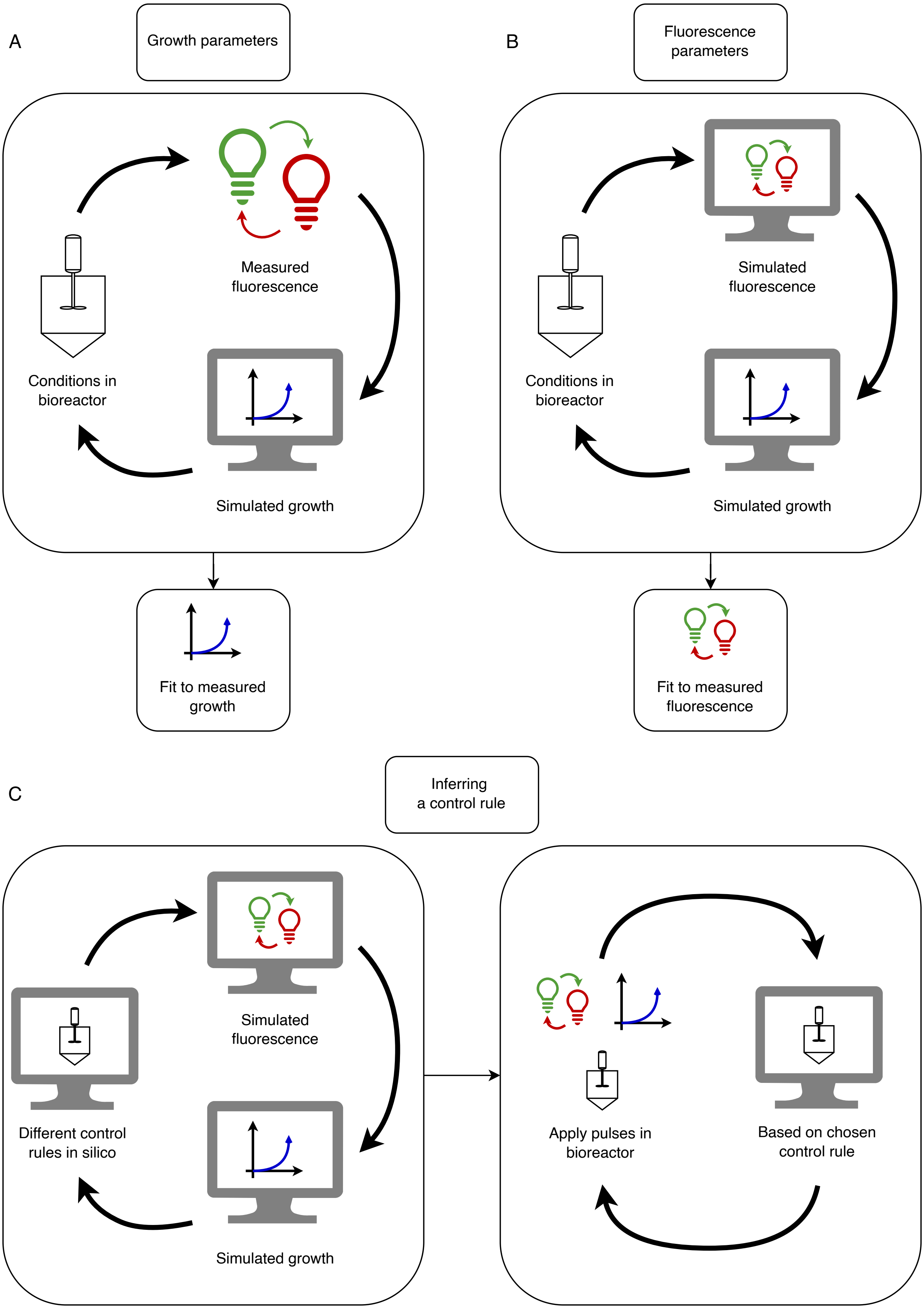
A
Growth parameters
Measured fluorescence
Conditions in bioreactor
Simulated growth
Fit to measured growth
B
Fluorescence parameters
Simulated fluorescence
Conditions in bioreactor
Simulated growth
Fit to measured fluorescence
C
Inferring a control rule
Simulated fluorescence
Different control rules in silico
Simulated growth
Apply pulses in bioreactor
Based on chosen control rule

Figure S7: Modular models make it possible to represent different parts of the process separately. **A** To estimate growth parameters, the conditions in the bioreactors and the fluorescence, measured or set during the process, were considered as input data. The model represented the growth of the population based on the measured phenotypes and was optimised using the measured biomass. **B** The fluorescence parameters are directly impacted by the concentration of inducer in the bioreactor, but the differences in growth rates between phenotypes also impacts their relative abundance. Based on the previously determined growth model, a fluorescence model was adjusted to represent reality. **C** Once this is done, it is possible to represent any experiment in silico, and possibly validate them in vivo.

For the non-growing phenotype, a maximum limit was applied to the growth rate with a simple threshold, considering that the limit in growth rate due to SerA auxotrophy should be independent from the glucose metabolism:

$$\mu_{i,l} = \min\left(\mu_i \ , \ \mu_{i,\mathrm{limited}}\right) \ , \quad \text{(SE 2)}$$

where $\mu_{i,\mathrm{l}}$ is the growth rate of the SerA-limited phenotype, $\mu_i$ is the growth rate of the growing phenotype (Equation SE 1), and $\mu_{i,limited}$ is the maximum growth rate of the limited phenotype.

In addition to these equations, to represent growth, the biomass yield $Y_{\frac{X}{S},i}$ (in $\mathrm{TCD_{biomass}}/\mathrm{g_{substrate}}$, to correspond to the measured biomass data) of each strain was considered identical for both phenotypes. This biomass yield was measured by considering the increase in biomass during batches, before the start of the continuous phase, where all 5 $\mathrm{g{\cdot}L^{-1}}$ of glucose provided are consumed.

The growth rates of each strain in bioreactor did not exactly match the measurements made in the Biolector. Therefore, they were re-calculated based on the pulse experiments (2024-08-08-TSGPulses1 and 2024-07-11-TSY2Pulses1), where, during the first series of pulses, biomass decreases, there is no mutation yet (as there is a biomass response), and the growth rate should be close to its maximum for both phenotypes. By considering the ratio of phenotypes as it is provided by fluorescence and the biomass and the observed derivative of the biomass, an approximation of the maximum growth rates in the continuous phase could be made (see Table 2).

Table 2: Maximum growth rates of each phenotype, determined based on Biolector and continuous bioreactor data.

| Strain | Phenotype | Growth rate in Biolector (in $\mathrm{h^{-1}}$) | Growth rate in bioreactor (in $\mathrm{h^{-1}}$) |
|---|---|---|---|
| Toggle_green | Growing | 0.53 | 0.75 |
| | Non-growing | 0.29 | 0.23 |
| Toggle_yellow | Growing | 0.44 | 0.66 |
| | Non-growing | 0.26 | 0.32 |

The last parameter, $K_{S,i}$, was determined based on glucose measurements during steady-state of single strains before any inducer was added.

Based on the measured parameters, the biomass curves could be reproduced given the fluorescence. This held true even when mutations appeared, as the fluorescence of mutated cells still indicated their ability to grow. For the population exhibiting no fluorescence at all at the end of the 2024-10-02-TSY2Pulses2 experiment, which would be counted as non-growing, the predicted overall biomass stayed stable since the non-growing population was small enough to be compensated by an increased growth of the growing phenotype. Notably, for Toggle_yellow, while we could reproduce the overall decrease in biomass, the experiments reached a stable biomass during the series of 15 ng·mL$^{-1}$ Tc pulses. This means that the assumption made to find the maximum growth rate (the growth rate of each phenotype is close to its maximum at this point) is not entirely valid. When running co-culture model, we adjusted the growth rate of Toggle_yellow to match the observed biomass decrease without inducer.

#### 2.3 Determination of induction parameters

Using the previous growth parameters, it is possible to isolate the behaviour of fluorescence in the bioreactor. For the Toggle_yellow strain, when tetracycline is added, a portion of the population switches to the non-growing phenotype. While we did make a model for this behaviour (see code at `https://gitlab.uliege.be/mipi/published-software/2025-syntheticniches`), it is never useful for the co-culture as Toggle_yellow grows more slowly than Toggle_green and is never induced in co-cultures. This model breaks similarly to Toggle_green when mutations appear.

For the Toggle_green strain, the behaviour of the median fluorescence can be reproduced using an inhibition model based on IPTG concentration:

$$\frac{\mathrm{dGFP}}{\mathrm{d}t} = k_{\mathrm{GFP}} \frac{K_{\mathrm{IPTG}}}{\mathrm{IPTG} + K_{\mathrm{IPTG}}} (\mathrm{GFP}_0 - \mathrm{GFP}) - \mu\, \mathrm{GFP}\ , \tag{SE 3}$$

where GFP is the fluorescence of the growing phenotype, associated to growth, $k_{\mathrm{GFP}}$ is the maximum production rate of GFP, $K_{\mathrm{IPTG}}$ is the IPTG concentration at half-maximum production rate, $\mathrm{GFP}_0$ is the maximum fluorescence theoretically possible, and $\mu$ is the growth rate of the growing phenotype.

These parameters could be determined based on the fluorescence data during the 2023-10-26-TSGPulseTest experiment in order to reproduce the fluorescence behaviour for the 2024-08-08-TSGPulses1 and 2024-09-05-TSGPulses2 experiments. Since the 2023-10-26-TSGPulseTest consists of pulses where the fluorescence has time to decrease and recover to base levels, it provides both the loss and recovery dynamics of GFP. In particular, the parameters could be determined as follows:

- If $\mathrm{GFP}_{\mathrm{max}}$ is the maximum measured GFP median, then the recovery part of the fluorescence, where IPTG is absent, can be expressed as:

$$\mathrm{GFP}_{\mathrm{max}} - \mathrm{GFP}(t) = c \exp\left(-\left(k_{\mathrm{GFP}} + \mu\right) t\right) \tag{SE 4}$$

  And the parameter $k$ can be inferred by fitting an exponential to the data.

- Then, based on the fluorescence at steady-state, the parameter $\mathrm{GFP}_0$ can be determined as:

$$\mathrm{GFP}_0 = \mathrm{GFP}_{\mathrm{max}} \frac{k_{\mathrm{GFP}} + \mu}{k_{\mathrm{GFP}}} \tag{SE 5}$$

- Finally, $K_{\text{IPTG}}$ can be determined by considering the fluorescence at the minimum median fluorescence of each pulse, where ${}^{\text{dGFP}}/_{\text{d}t} = 0$:

$$0 = k_{\text{GFP}} \frac{K_{\text{IPTG}}}{\text{IPTG}_{\text{min}} + K_{\text{IPTG}}} (\text{GFP}_0 - \text{GFP}_{\text{min}}) - \mu \, \text{GFP}_{\text{min}} \, , \tag{SE 6}$$

leading to:

$$k_{\text{apparent}} = \mu \frac{\text{GFP}_{\text{min}}}{\text{GFP}_0 - \text{GFP}_{\text{min}}} \tag{SE 7}$$

$$= k_{\text{GFP}} \frac{K_{\text{IPTG}}}{\text{IPTG}_{\text{min}} + K_{\text{IPTG}}} \tag{SE 8}$$

$$K_{\text{IPTG}} = \frac{k_{\text{apparent}}}{k_{\text{GFP}} - k_{\text{apparent}}} \text{IPTG}_{\text{min}} \tag{SE 9}$$

This provides the required data to reproduce the fluorescence median, but in order to use that fluorescence and to determine the growth rate, there needs to be a relationship between the median and the percentage of growing cells. As an approximation, we decided to consider that the increase in non-growing cells was a direct function of the median, such that if $X_1$ and $X_2$ are the growing and non-growing biomasses respectively, then:

$$\frac{\text{d}X_1}{\text{d}t} = (\mu_{\text{TSG}} - D) X_1 - X_1 \exp\left(-\frac{\text{GFP}}{\tau}\right) \tag{SE 10}$$

$$\frac{\text{d}X_2}{\text{d}t} = (\mu_{\text{TSG},l} - D) X_2 + X_1 \exp\left(-\frac{\text{GFP}}{\tau}\right) \tag{SE 11}$$

(SE 12)

when the median is decreasing. It can be determined from a linear regression of experimental data during the decrease in GFP as everything in EQUATION SE 11 can be extracted from biomass and fluorescence data except for $\tau$. The resulting parameters work surprisingly well to represent other experiments despite the simplicity of the model, as long as there is no mutation.

#### 2.4 Co-culture model

With both models established, it is possible to combine them to represent a co-culture of TOGGLE_green and TOGGLE_yellow. A co-culture without any inducer pulses closely reproduces the the observed behaviour, provided we adjust the growth rate of TOGGLE_yellow, as mentioned in section 2.2. Then, by applying pulses in silico, the model indicates that the TOGGLE_yellow strain can be maintained in the medium, provided there is no mutation in the culture (see FIGURE S8 for the control rule that was applied in practice). As experiments showed, mutations make this model invalid very quickly, despite applying at most 2 pulses in 7 h and 3 pulses in 20 h.

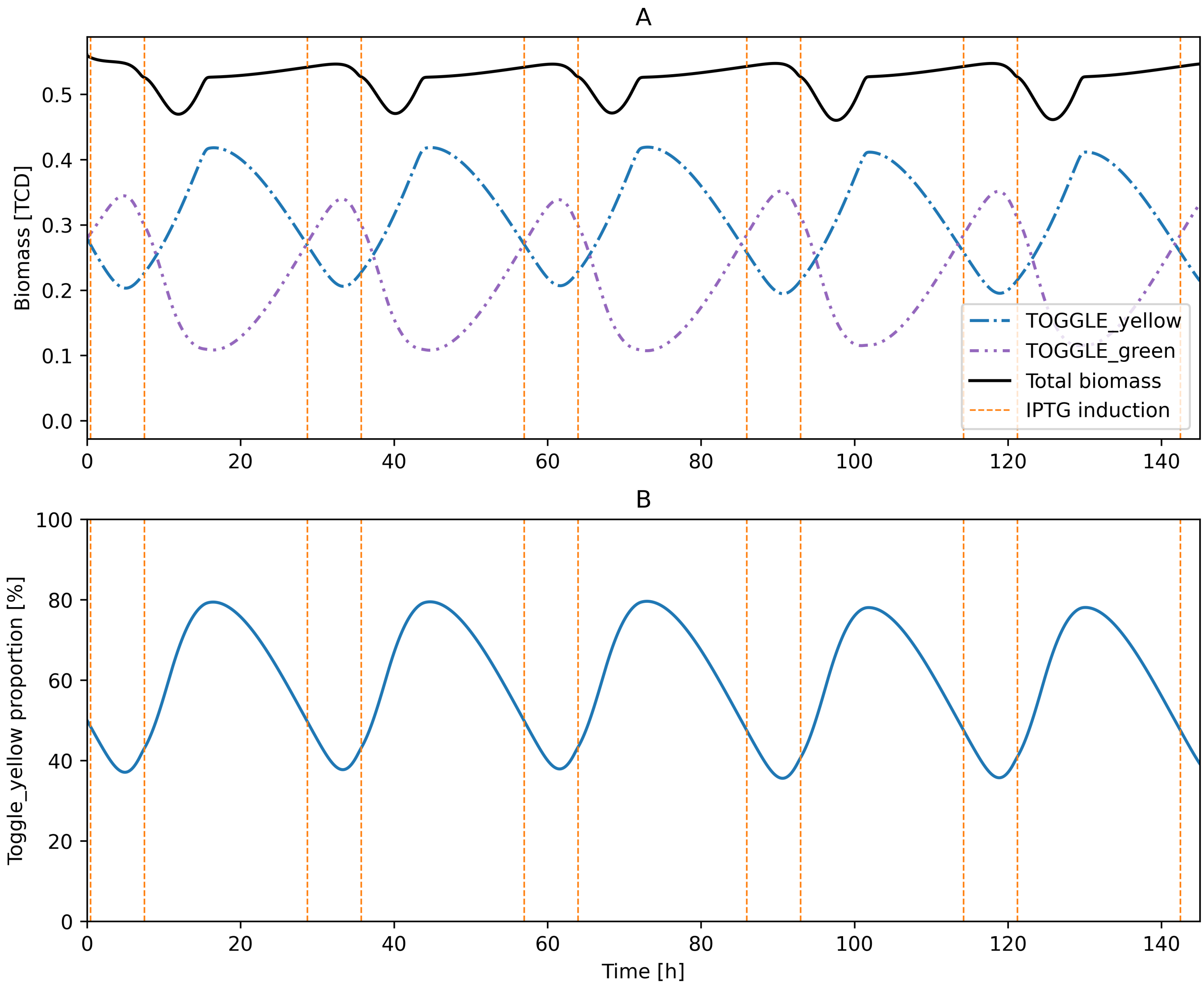

Figure S8: **A** Predicted biomass behaviour in continuous culture when the control rule is applied. **B** Corresponding percentage of Toggle_yellow in the co-culture.

### 3 Supplementary file list

The supplementary files available at `doi.org/10.5281/zenodo.17663997` contain the following documents:

- `genetic_constructs.7z` contains the sequences obtained for the TOGGLE_green and TOGGLE_yellow plasmids, as well as the sequences for the modified genomic DNA regions $\Delta$*lacAYZI* and $\Delta$*serA* in *E.coli* MG1655.
- `cytometer_calibration.7z` contains the fcs files used for calibration of fluorophores and differences between cytometers.
- Each `experiment_*.7z` file contains the corresponding experiment data, as well as associated notes and plots relevant to understand it.
- `mut_sequences.7z` contains the sequences obtained when checking for mutations.

All the relevant code for data analysis is available at `https://gitlab.uliege.be/mipi/published-software/2025-syntheticniches`.
